# Systematic study of hybrid triplex topology and stability suggests a general triplex-mediated regulatory mechanism

**DOI:** 10.1101/2024.05.28.596189

**Authors:** Vito Genna, Guillem Portella, Alba Sala, Montserrat Terrazas, Núria Villegas, Lidia Mateo, Chiara Castellazzi, Mireia Labrador, Anna Aviño, Adam Hospital, Albert Gandioso, Patrick Aloy, Isabelle Brun-Heath, Carlos Gonzalez, Ramon Eritja, Modesto Orozco

## Abstract

By combining *in-silico*, biophysical and *in-cellulo* experiments, we decipher the topology, physical and potential biological properties of hybrid-parallel nucleic acids triplexes; an elusive structure at the basis of life. We found that hybrid triplex topology follows a stability order: r(Py)-d(Pu)·r(Py)> r(Py)-d(Pu)·d(Py)> d(Py)-d(Pu)·d(Py)> d(Py)-d(Pu)·r(Py). The r(Py)-d(Pu)·d(Py) triplex is expected to be the preferred in the cell as it avoids the need to open the duplex reducing the torsional stress required for triplex formation in the r(Py)-d(Pu)·r(Py) topology. Upon a massive collection of melting data, we have created the first predictor for hybrid triplex stability. Leveraging this predictor, we conducted a comprehensive scan to assess the likelihood of the human genome and transcriptome to engage in triplex formation. Our findings unveil a remarkable inclination - of both the human genome and transcriptome - to generate hybrid triplex formation, particularly within untranslated (UTRs) and regulatory regions, thereby corroborating the existence of a triplex-mediated regulatory mechanism. Furthermore, we found a correlation between nucleosome linkers and TFS which agree with a putative role of triplexes in arranging chromatin structure and local/global level.

## INTRODUCTION

Triplexes are formed when a poly-purine segment of a duplex is recognized by a third oligonucleotide strand (the TFO; triplex forming oligonucleotide) by means of specific hydrogen bond interactions along the major groove ^1–5^. The TFO can be arranged parallel or antiparallel to the purine (Pu) strand. The triads (T-A·T, C^+^-G·C and G-G·C) present in parallel triplexes are stabilized by means of Hoogsteen hydrogen bonds, while reverse Hoogsteen hydrogen bond pattern stabilizes triads (A-A·T, G-G·C and T-A·T) in antiparallel triplexes (where “-” refers to Hoogsteen/reverse Hoogsteen and “·” refers to Watson-Crick pairings). Isosteric consideration favors triplexes where the third strand is either homopyrimidine (parallel triplexes; pyrimidine (Py) motif) or homopurine (antiparallel triplexes; purine (Pu) motif) ^4–6^. Despite the pH dependence of the C^+^-G·C triad, the parallel triplexes are more stable than the anti-parallel ones under physiological conditions ^6–9^.

Early fiber diffraction models suggest an A-type conformation for the DNA triplex^10^, but several NMR experiments and exhaustive molecular dynamics (MD) simulations demonstrated that the DNA triplex shows a “B-like” conformation, with sugars in the South conformation and triads perpendicular to the helix axis ^11–18^. The duplex major groove (MG) is divided by the TFO in two grooves^14,17,18^: one very narrow with purine C8 in the bottom (mMG in Figure 1), and the other very wide (MMG in Figure 1), covering all the region between the TFO and the third strand of the duplex (Figure 1). The presence of the TFO blocks the major-groove recognition pattern between transcription factors and the DNA duplex, and while the MMG can be recognized by some proteins ^18–20^ the general and main effect of triplex formation is the inactivation of DNA transcription; the effect being maximized if the triplex is formed in the regulatory regions ^4,21–24^. Very interestingly ^25,26^, promoters in most organisms, including humans, are highly enriched in poly-Pu sequences (triplex target sequences; TTS), suggesting that a large number of genes could be inactivated by triplex formation ^25^ if a suitable TFO is available.

**Figure 1.**
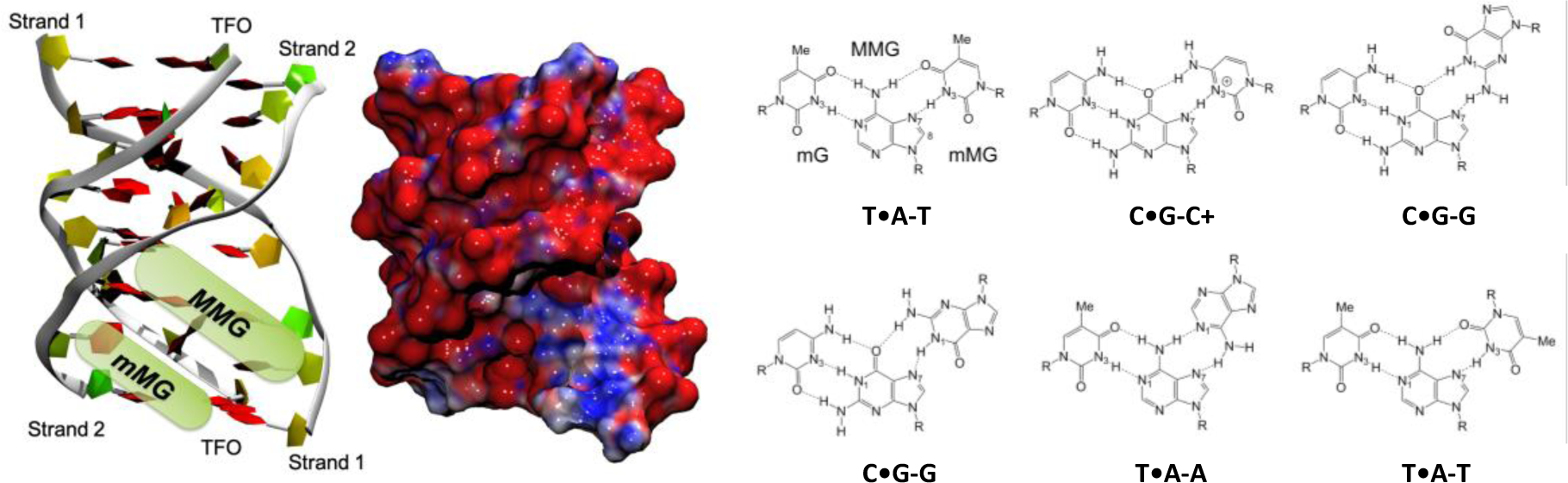
Left panel: tridimensional representation of a triplex with the two sections of the Major Groove highlighted. Right panel. Schematic representation of T·A-T, C·G-C^+^, C·G-G, C·G-G, T·A-A and T·A-T triads in parallel (top row) and antiparallel (bottom row) triplexes.

The possibility to block DNA expression by adding a DNA-based TFO has been exploited in the so-called anti-gene therapy ^21–24,27–38^. However, in normal cellular conditions, single stranded DNA are rare, and putative TFOs are RNA sequences, which act in some cases as regulators of gene expression ^39–45^. These experimental findings, combined with bioinformatic analysis ^25,26^ suggest the existence of an ancient feedback regulatory mechanism based on the formation of triplexes with TTS in a regulatory gene and TFO defined by the RNA of a regulated gene or an accessory regulatory RNA. Very recently, bioinformatics data have been published showing a significant correlation between human HiC contact maps and regions that can form triplexes with a third strand made of long non-coding RNAs ^46,47^ suggesting a correlation between triplex formation and genome structural organization ^46,47^.

Understanding the role of hybrid triplexes in gene regulation and chromatin structure first requires a good knowledge of their structural characteristics. Unfortunately, contrary to pure DNA triplexes ^1–18,48–51^, little and contradictory information exists on the structure of hybrid triplexes ^48–52^. We combine here MD simulations and biophysical experiments to explore the stoichiometry, topology, stability, structure and dynamics of hybrid triplexes. We found, both theoretically and experimentally, a general order of stability: r(Py)-d(Pu)·r(Py)> r(Py)-d(Pu)·d(Py)> d(Py)-d(Pu)·d(Py)> d(Py)-d(Pu)·r(Py), the rest showing little stability, except when the d(Pu) is made of a poly-d(A), in which case the ordering is d(Py)-r(Pu)·r(Py)> d(Py)-d(Pu)·d(Py)> r(Py)-r(Pu)·r(Py)> r(Py)-r(Pu)·d(Py), with little structural differences between the stable hybrid triplexes. We centered our attention in the triplex that is likely to be more prevalent in cellular conditions: r(Py)-d(Pu)·d(Py), developing and validating a stability predictor which allows us to scan for the stability of these triplexes under a range of conditions. Applying this predictor to genomic and transcriptomic data, the likelihood of hybrid triplex formation in human cells is analyzed. A large prevalence of these triplexes is found, being very abundant in regulatory regions (promoters and 5’UTR) and involving mainly miRNAs as TFOs. These findings provide strong support to the hypothesis of an ancient RNA-based triplex-mediated regulatory mechanism. Furthermore, triplexes are located at positions where they can help to fix chromatin structure, both locally and globally.

## RESULTS AND DISCUSSION

### Stability of Homo-polymers and Hybrid Triplexes

Melting experiments were first performed using different homopyrimidine triplexes as TFO and homopurine-homopyrimidine hairpins as TTSs. The use of hairpins (polyethylene glycol was used as loop) has the advantage to minimize the formation of other competing structures in the TTS, such as reverse Watson-Crick ^53,54^, Hoogsteen duplexes ^55–59^, quadruplexes and others ^60–62^ which will introduce noise in the T_m_ estimates). We consider 3 compositions of the hairpin: (100% A·T/U (**I-IV**); 70% A·T/U (**V-VIII**) and 50% A·T/U (**IX-XII**)), we do not consider higher percentages of guanines as this will increase the risk of quadruplex formation. With these compositions we create all the combinations of DNA and RNA in the hairpin: d(Pu)·d(Py) (**I**, **V** and **IX**); r(Pu)·d(Py) (**II**, **VI** and **X**); d(Pu)·r(Py) (**III**, **VII** and **XI**) and r(Pu)·r(Py) (**IV**, **VIII** and **XII**)) and incubate them with the corresponding homopyrimidine TFO (DNA with 100%, 70% or 50% T (**1**, **3** and **5**); RNA with 100%, 70% or 50% U (**2**, **4** and **6**). Combination of all TFOs with all TTS leads to 24 potential triplexes whose stability was measured by the corresponding melting curves (recorded in all cases at pH 6.0; see Methods). Results (Figure 2) show melting temperatures in the range T<15°C (not detectable) to 52 °C. Triplexes with 100% A·T/U show in general a poor stability with a decreasing order of stability d(Py)-r(Pu)·r(Py)> d(Py)-d(Pu)·d(Py)> r(Py)-r(Pu)·r(Py)> r(Py)-r(Pu)·d(Py) the rest being not detectable (Figure 2). When the ratio of G·C increases the triplexes become more stable, showing a quite well-defined order of stability (Figure 2): r(Py)-d(Pu)·r(Py)> r(Py)-d(Pu)·d(Py)> d(Py)-d(Pu)·d(Py)> d(Py)-d(Pu)·r(Py)> r(Py)-r(Pu)·d(Py) ≈ r(Py)-r(Pu)·r(Py). At the studied pH, the increase in the ratio G·C/A·T implies an increase in the stability of the triplex. For a given G·C/A·T ratio the two most stable triplexes are those with RNA in the TFO with d(Pu)·r(Py) preferred over d(Pu)·d(Py) in the TTS.

**Figure 2.**
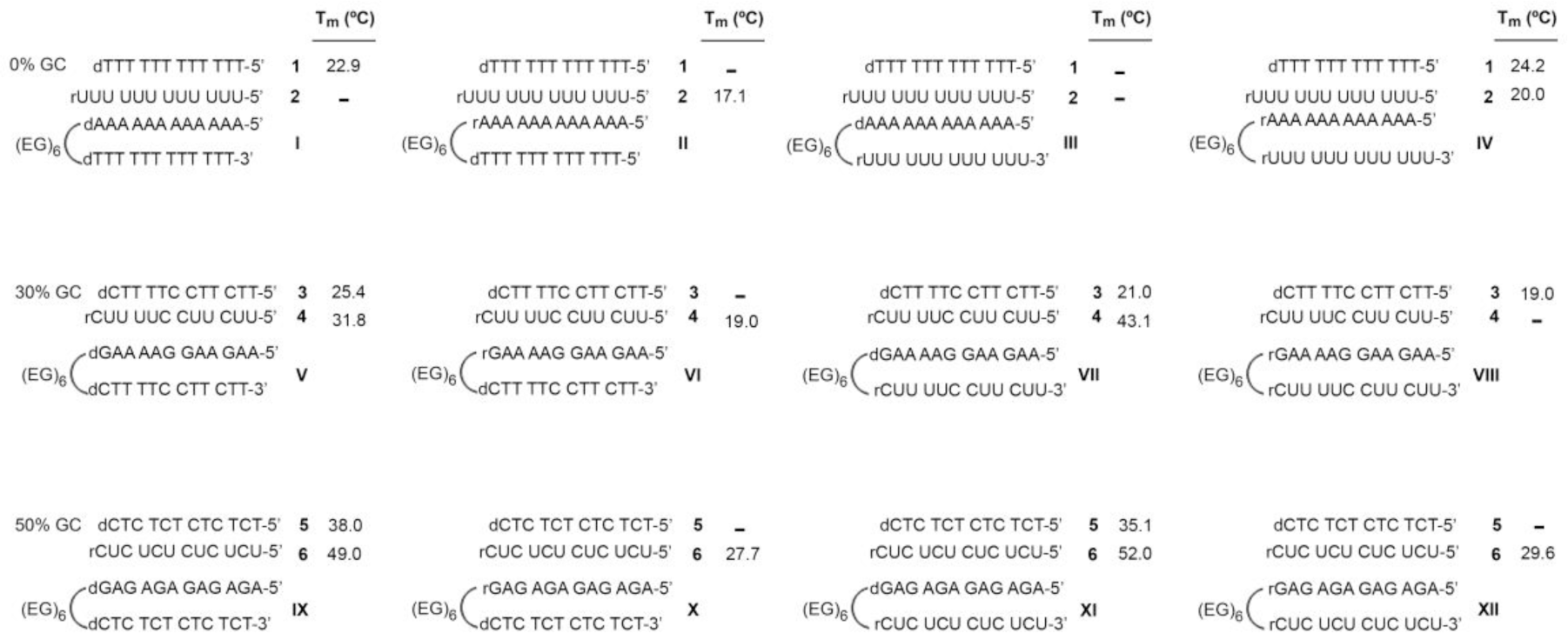
Melting temperatures of d(Py)-d(Pu)·d(Py), r(Py)-d(Pu)·d(Py), d(Py)-r(Pu)·d(Py), r(Py)-r(Pu)·d(Py), d(Py)-d(Pu)·r(Py), r(Py)-d(Pu)·r(Py), d(Py)-r(Pu)·r(Py) and r(Py)-r(Pu)·r(Py) 12mer triplexes of 100%, 70% and 50% A·T/U content in 10 mM sodium cacodylate buffer (pH 6.0) containing 100 mM NaCl and 10 mM MgCl_2_ (see Materials and Methods for details). Only the T_m_ of the triplex transition is indicated in all cases. For results at higher pH see below.

In order to confirm that melting experiments were really analyzing triplex**→**duplex transitions instead of other processes related to strand invasion with R-loop formation, we repeated melting experiments for triplexes d(Py)-d(Pu)·d(Py), r(Py)-d(Pu)·d(Py), d(Py)-d(Pu)·r(Py) and r(Py)-d(Pu)·r(Py) for the case of 70% A·T/U in the TTS) at higher pH values (from 6.0 to 8.0; see Figure 3a-h) finding a reduction of T_m_, consistent with triplex formation. The triplex nature of the structures was further confirmed by means of ^1^H NMR spectroscopy. Thus, ^1^H-NMR spectra of hairpin **VII** and its equimolar mix with **4** different temperatures show the expected decay of imino signal intensities as temperature increases (panels i and j of Figure 3). In the spectra of **VII**, four guanine imino signals involved in WC base pairs (12-13 ppm) are observed at T = 5°C, being two of them still observed at 45°C. Most probably, they correspond with the two central GC base-pairs. In addition, uracil imino signals forming Watson-Crick AU base-pairs are observed around 14 ppm. These NMR data are fully consistent with formation of a hairpin structure as shown in the insert of the Figure 3i. Upon addition of triplex forming RNA **4**, the number of exchangeable proton signals increases drastically. Formation of GC Hoogsteen base pairs is demonstrated by the observation of protonated cytosine imino signals around 15 ppm and their corresponding amino signals around 10 ppm. Additional sharp and well-dispersed imino signals can be observed in the 13 to 14 ppm region, consistent with the formation of additional AU base pairs. Most of these signals are observed at 45°C indicating the formation of a very stable structure (Figure 3j). Overall, the NMR spectra of the mix are consistent with formation of the expected parallel triplexes (see Figure 1). Overall, our experiments explain apparently contradictory previous data, agreeing with Crother’s estimates^51^ obtained for 66% GC triplexes and with Dervan’s data ^51,52^ collected for 81% AT triplexes.

**Figure 3.**
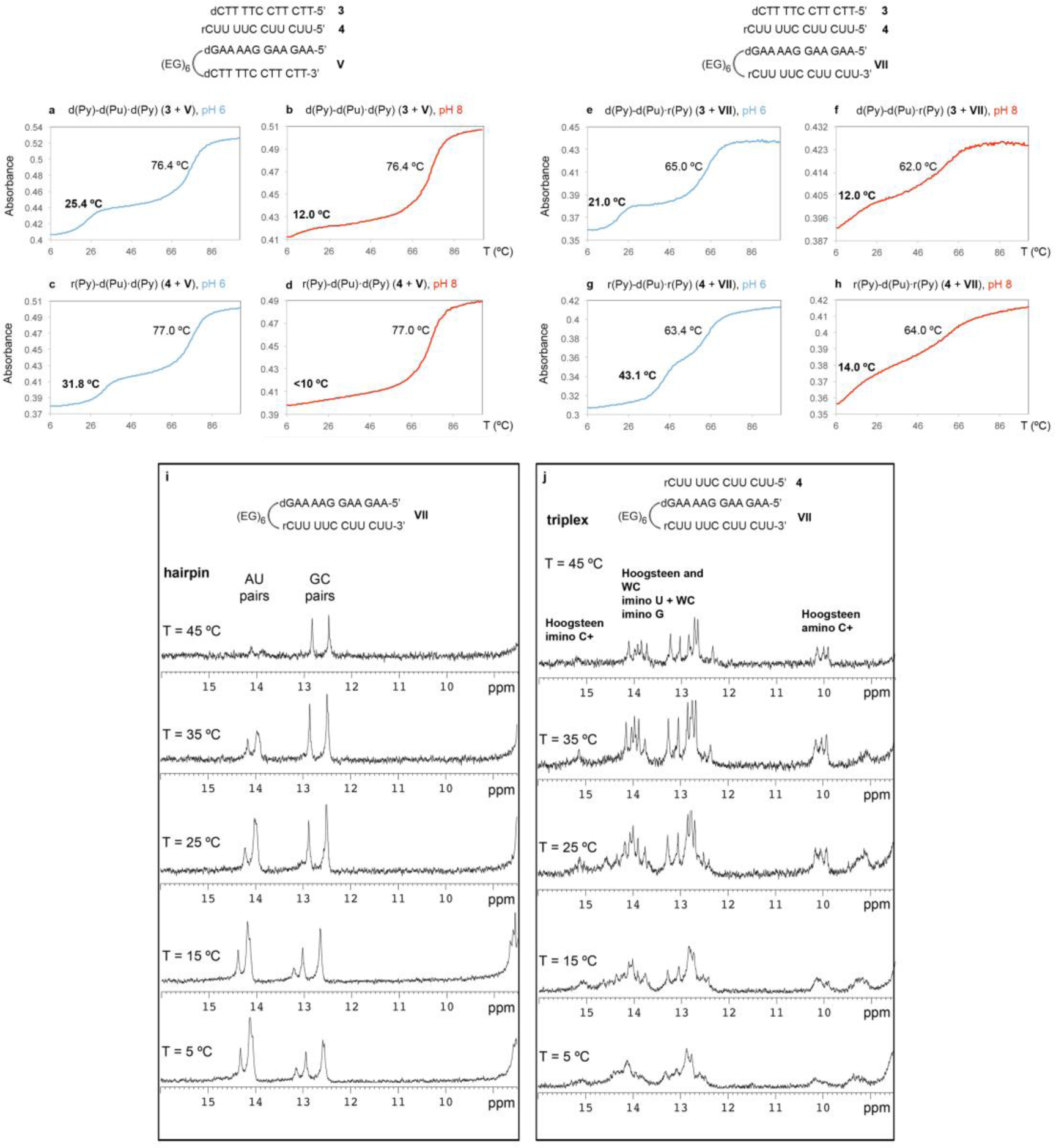
**a-h**. Thermal stability of d(Py)-d(Pu)·d(Py) (**3** + **V**), r(Py)-d(Pu)·d(Py) (**4** + **V**), d(Py)-d(Pu)·r(Py) (**3** + **VII**) and r(Py)-d(Pu)·r(Py) (**4** + **VII**) 12mer triplexes in 10 mM sodium cacodylate buffer (pH 6.0) containing 100 mM NaCl and 10 mM MgCl_2_ (**a, c, e, g**) and in 10 mM sodium cacodylate buffer (pH 8.0) containing 100 mM NaCl and 10 mM MgCl_2_ (**b, d, f, h**). Melting temperatures (T_m_) for the duplex and triplex are indicated in each case (in bold: T_m_ corresponding to the triplex). **(i, j**) ^1^H NMR spectra of the imino region of D·R hairpin **VII** (**i**) and r(Py)-d(Pu)·r(Py) triplex **4** + **VII (j)** acquired at 5 °C, 15 °C, 25 °C, 35 °C and 45 °C in 30 mM phosphate buffer (pH 6.0) containing 100 mM NaCl and 10 mM MgCl_2_.

Very interestingly, hybrid triplexes d(Py)-d(Pu)·r(Py) are quite unstable compared to the r(Py)-d(Pu)·d(Py) ones, which suggests that triplexes with a 2:1 (DNA:RNA) stoichiometry show a topology r(Py)-d(Pu)·d(Py). Furthermore, triplexes r(Py)-d(Pu)·r(Py) which are intrinsically very stable, are expected to be disfavored in the cell, as its formation requires strand invasion, which would generate a strong topological stress in the DNA duplex and an unstable unpaired d(Py) strand. So, present results suggest that r(Py)-d(Pu)·d(Py), with TTS being the genomic DNA and the TFO being expressed RNAs will be the most stable triplex topology in the cell.

### The Structure and Dynamics of the Hybrid Triplexes

To gain structural and mechanistic insights on the structure and stability of hybrid triplexes, we performed a set of extensive molecular dynamics (MD) simulations (see Methods) of 6 triplexes in aqueous solution: r(Py)-d(Pu)·r(Py), r(Py)-d(Pu)·d(Py), d(Py)-d(Pu)·d(Py), d(Py)-d(Pu)·r(Py), r(Py)-r(Pu)·r(Py) and r(Py)-r(Pu)·d(Py). Hydrogen bonds (H-bonds) are not equally conserved in all triplexes, the differences being especially remarkable for the Hoogsteen ones, which agree with the experimental difference in stability between them. Thus, for the stable r(Py)-d(Pu)·r(Py) and r(Py)-d(Pu)·d(Py) triplexes (Figure 2), around 96% of Watson-Crick and 95% of Hoogsteen hydrogen bonds are preserved, while for “low-stability” r(Py)-r(Pu)·r(Py) and r(Py)-r(Pu)·d(Py) triplexes massive disruption of H-bonds is found, leading to structural disruption (Figure 4A). In fact, looking at the conservation of H-bonding we can define a “theoretical stability ordering”: r(Py)-d(Pu)·r(Py)≥ r(Py)-d(Pu)·d(Py)> d(Py)-d(Pu)·d(Py) ≥ d(Py)-d(Pu)·r(Py)> d(Py)-d(Pu)·r(Py) ≈ r(Py)-r(Pu)·r(Py), which matches the experimental one (Figure 2). RMSd distributions (Figure 4B) confirm the stability ranking derived from H-bond analysis. Overall, agreement between theoretical and experimental estimates provide very strong confidence on the reliability of the atomistic MD simulations ^63–66^. Analyzing the different sampled structures, we found that all hybrid triplexes show similar structures, which are not far from the homopolymeric ones, in terms of groove geometries and helical coordinates (see Figure 4; see Suppl. Table S1 and references ^10–17,63^). There are however interesting differences depending on the stoichiometry and topology of the hybrid triplexes. For example, 2(RNA):1(DNA) triplexes are in general more “A-like” as reflected in lower twist-higher roll than 1(RNA):2(DNA) triplexes (see Suppl. Table S1). As expected from previous studies on DNA·RNA duplexes ^63^ RNA-strands maintain N-puckering, while DNA strands sample South and East regions (Suppl. Table S2). The situation does not change much depending on the placement of the nucleic acid strand in the Watson-Crick or Hoogsteen strands.

**Figure 4.**
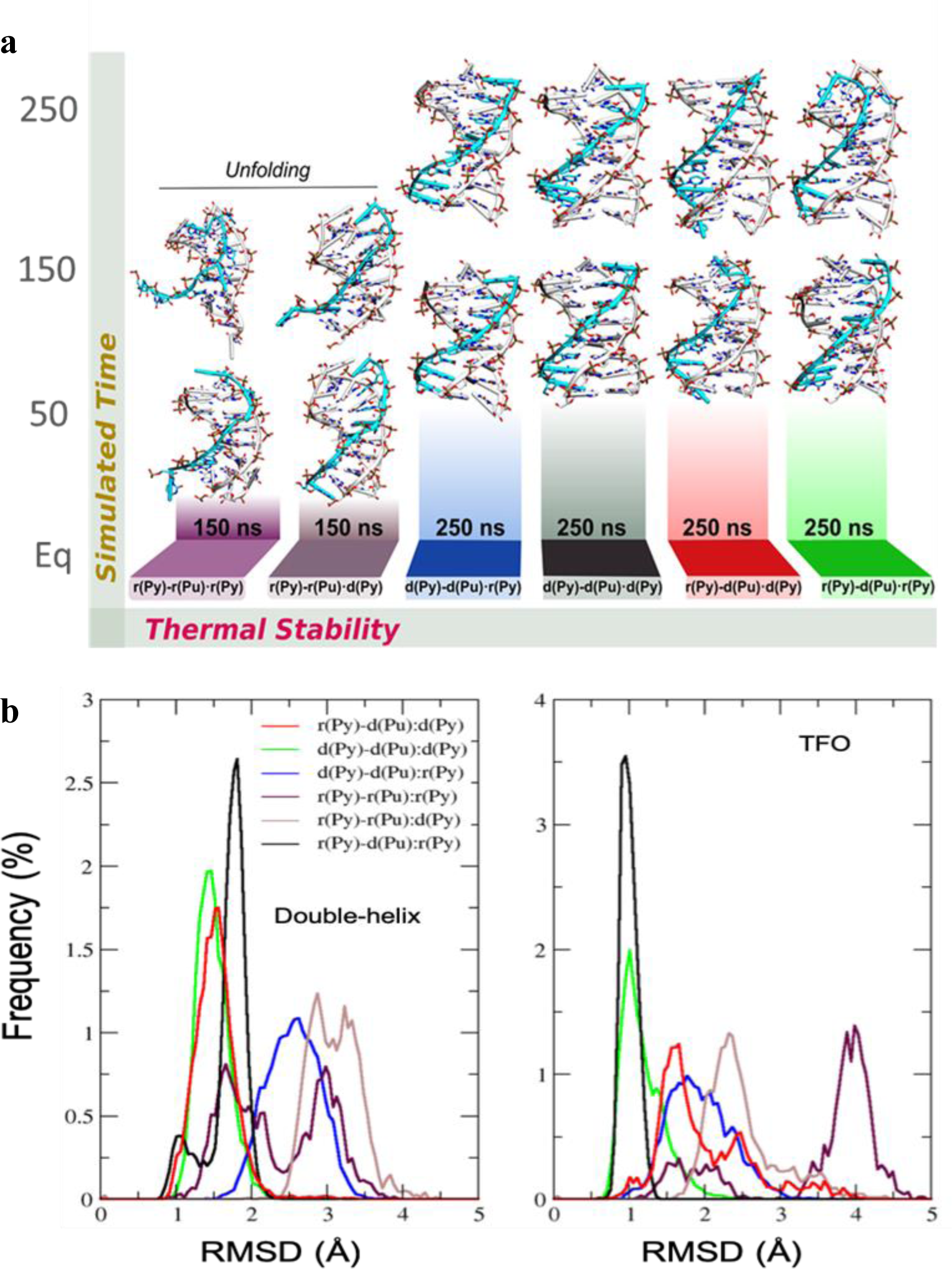
Selected results of the MD simulation of different hybrid triplexes. **a.** Evolution of the structure of the different triplexes along trajectories; **b.** Histograms of the RMSd in the TTS (left) and TFO (right) along the trajectories.

### Development and Validation of a Predictor of the Stability of r(Py)-d(Pu)·d(Py) Triplexes

As discussed above, the most stable triplex (r(Py)-d(Pu)·r(Py)) is not expected to have a large prevalence in the cell out of R-loop constructs. However, the second most stable triplex: r(Py)-d(Pu)·d(Py) can be easily formed by pairing an RNA segment with genomic DNA. Following Robert and Crothers approach ^50^ we trained a simple nearest neighbor model for r(Py)-d(Pu)·d(Py) to reproduce experimental data in a variety of triplexes (see Methods) in a variety of conditions. The refined method predicts experimental melting observables with root mean square errors around: 4.8 degrees (T_m_), and 0.7 kcal/mol (melting free energy), improving dramatically the accuracy obtained by transferring Roberts-Crothers DNA triplex method (see Figure 5A). Our predictions also outperform the widely used Triplexator software ^67^ which is unable to detect all stable triplexes at a given temperature; see Figure 5B.

**Figure 5.**
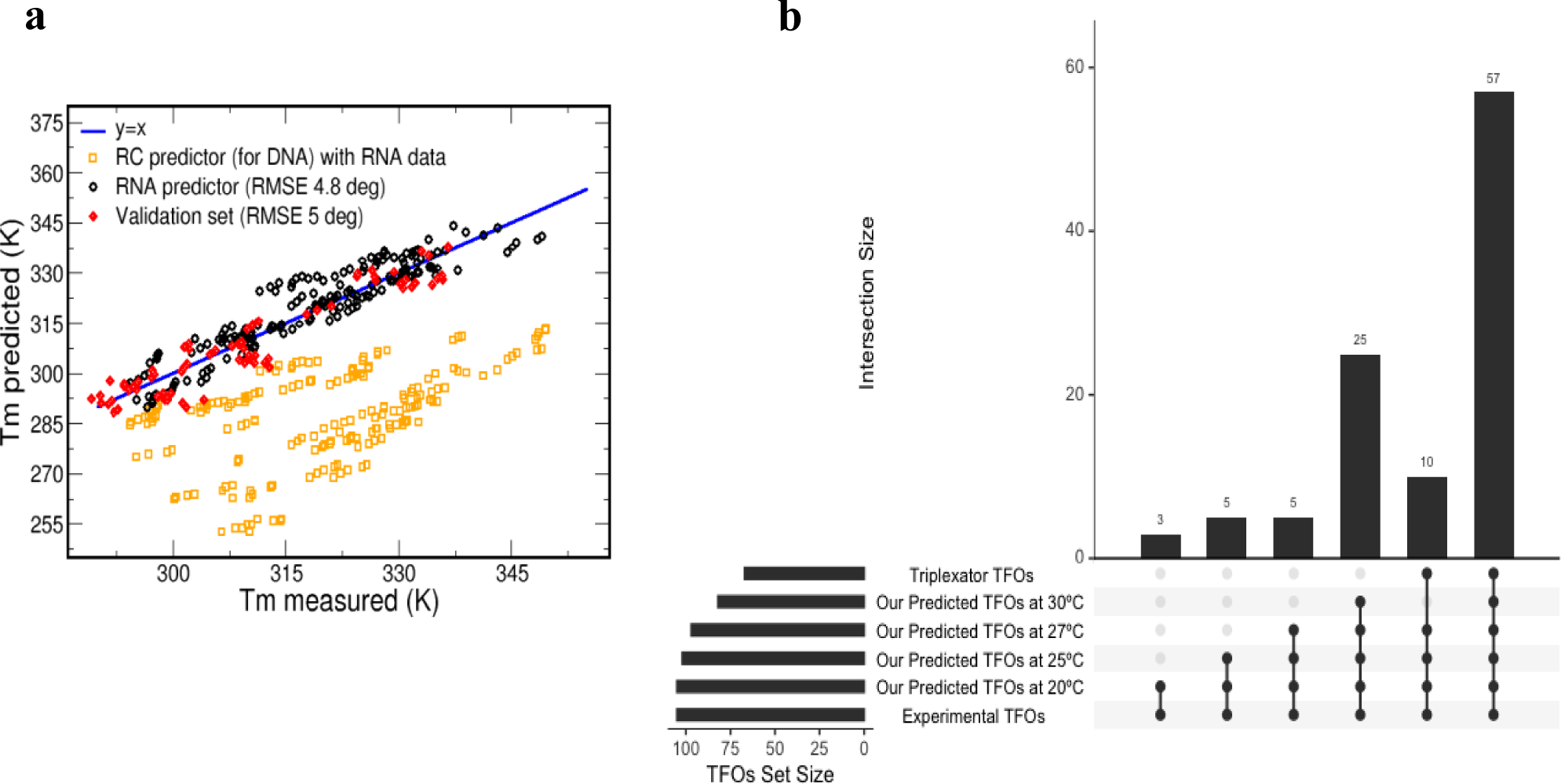
**a.** Predicted vs measured melting temperatures of triplexes. Values in yellow correspond to estimates obtained using Roberts-Crothers method for triplexes. Values in black to the data for the training set and that in red to data for a completely independent validation set (see Methods for details). **b.** Predicted TFOs from our model in comparison to Triplexator in the evaluation of stable triplexes at various temperatures. Results are shown as an intersecting upset plot.

To further validate the predictive power of our model, we designed a 50 nt polypyrimidine TFO (TFO **7**; Figure 6a) which, according to our method, should form stable triplexes (T_m_ values = 57 °C at pH 6.5 and 48 °C at pH 7.0) in the promoter region of the BRD7 gene. As shown in Figure 6a, synthetic TFO 7 interacts with a radiolabeled synthetic double-stranded DNA hairpin (**XIII**) comprising the target BRD7 polypurine sequence, forming a low-mobility complex. To validate the triplex nature of this complex, chromatin extracted from HeLa cells (see Materials and Methods for details) was treated with DNase I and proteinase K and sonicated to yield fragments of 200-300 nt (see Suppl. Figure S3). Two aliquots of this purified DNA were incubated with TFO 8, a biotinylated version of TFO 7, at pH 5.5 and pH 7.0 in the presence of RNase H to digest putative R-loops. The streptavidin-retained DNA was eluted and identified by qPCR amplification, finding significant DNA recovery when using BRD7 promoter-specific primers amplifying a 92 nt region just around the target triplex region (92Nt primers; Figure 6b; red panel, Suppl. Figure S4 and Suppl. Table S3). Interestingly, a decrease in the pH led to an increase in DNA recovery, which is consistent with pH-dependent stability in C-C·G triplex formation as captured by our predictor. Similar results were obtained when using promoter-specific primers amplifying a larger region (217 nt) around the target triplex region (217Nt primers; Figure 6b; yellow panel). On the contrary, no recovery was observed when using intronic-specific primers (Figure 6b; green panel), or when genomic DNA fragments were incubated with a TFO lacking the sequence matching the target region (TFO **9**), confirming the specificity of the above-described results and the triplex nature of the complex predicted by our model.

**Figure 6.**
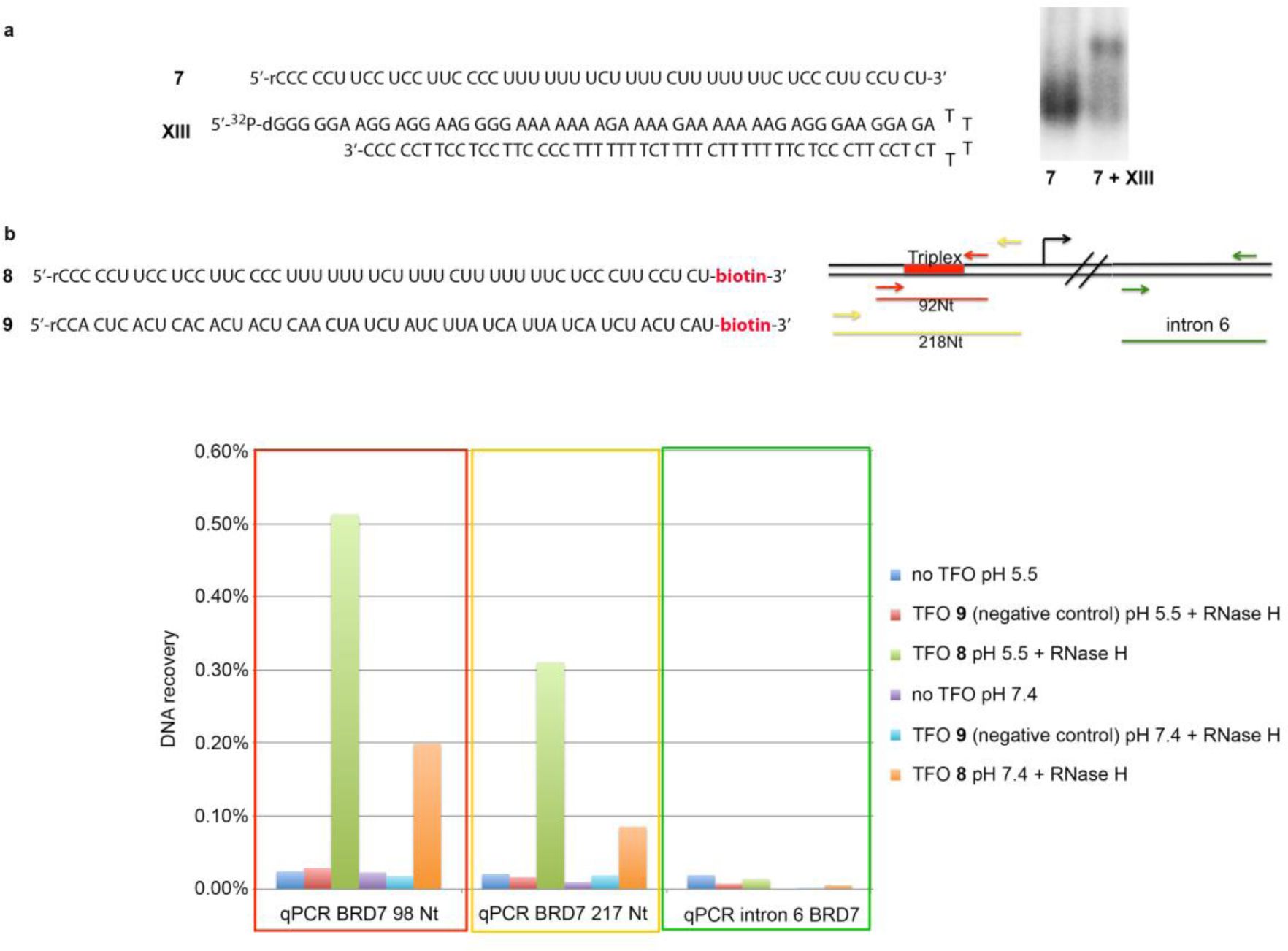
**a.** Electrophoretic mobility shift assays to analyze triplex formation between TFO **7** and hairpin **XIII**. **b.** Biotinylated TFOs used in this study (**8**: TFO targeting the polypurine A/G) site of BRD7 promoter region; **9**: negative control). Schematic depicting the position of the target triplex forming region and the primers used for DNA amplification. Upon binding to streptavidin beads, associated DNA was analyzed by qPCR using promoter-specific [92 NT (in red) or 217 NT (in yellow)] or intronic-specific (in green) primers.

### Formation of r(Py)-d(Pu)·d(Py) Triplexes in Human Cells

We used our predictor to screen for potential TFOs amongst annotated human lncRNAs and miRNAs from the gencode and miRbase ^68,69^ databases respectively (see Methods). We found a strong enrichment of TFO candidates (triplex Tm>30°) in both dataset in comparison to the population of expected TTSs from a random distribution (see random model in Methods) (Figure 7), suggesting potential r(Py)-d(Pu)·d(Py) triplex formation *in vivo*. When compared with the DNA-associated RNA isolated by Grummt and coworkers ^45^, we found that 44% or our predicted TFOs from lncRNAs and 51% from miRNAs were indeed found associated with DNA in a triplex structure. Both sets of TTS (from miRNA and lncRNA TFOs) were mapped to the human genome, and we observed an over-representation in promoters when analyzing stable triplexes (Tm > 30°), and in 5’UTR (in the case of miRNA’s TFO) when analyzing very stable Triplexes (Tm>45°C), suggesting that parallel r(Py)-d(Pu)·d(Py) triplexes form preferentially in regions important for the control of gene expression.

**Figure 7.**
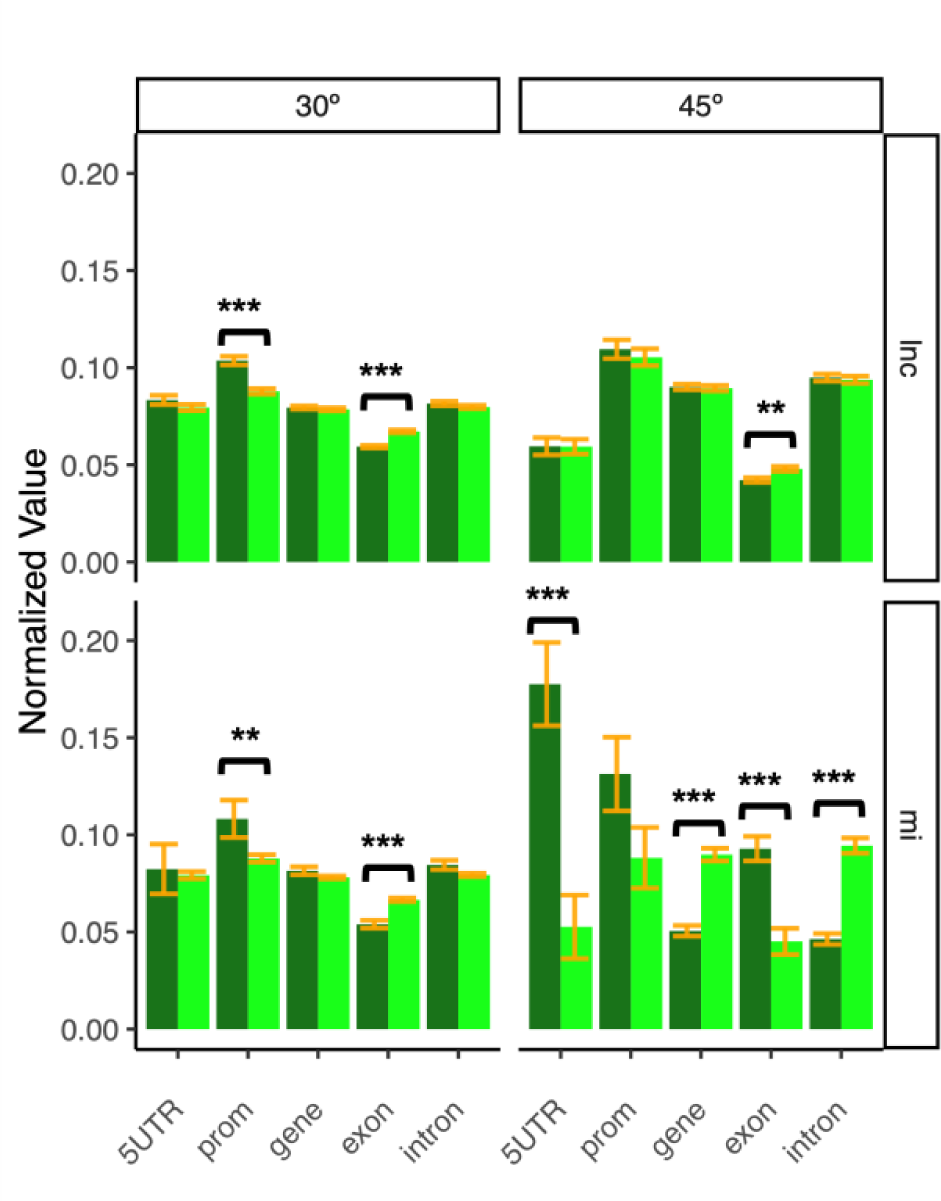
Overall representation across different genomic classes in lncRNAs (upper plots – dark green) and miRNAs (lower plots – dark green) against a random background model (light green), with the corresponding standard error bars (orange). RNA-DNA·DNA triplexes are shown as stable at 30° (left-most plots) or 45 °C (right-most plots) in the different regions. When a p-value is less than 0.05, it is flagged with one star (*). If a p-value is less than or equal to 0.01, it is flagged with two stars (**). If a p-value is less than or equal to 0.001 it is flagged with three stars (***).

GO analysis of the genes potentially controlled by RNA-DNA·DNA triplex formation with miRNAs and lncRNAs showed that these genes are frequently related to complex processes, such as development (see Suppl. Figures S5), with very significant hits in the development of the nervous system. It is tempting to speculate that stable triplexes generated by the binding of transcribed RNAs with genomic DNA can be involved in a fine-tuning regulatory mechanism, which was inherited from an ancient triplex-mediated DNA-RNA regulatory network. Note that this finding agrees well with the work from Pasquier et al. in Drosophila that showed that the genes targeted by TFOs were involved in development and morphogenesis ^70^.

### Final comment on role in chromatin structure

To investigate triplex formation in the context of chromatin, we predicted the putative TFOs from lncRNA and miRNAs expressed in lymphoblastic cells ^71^ and compared the location of their target sites along the human genome with a genome-wide map of nucleosome occupancy in human lymphoblastoid cell line ^72^. We observed a local minimum which coincides with the nucleosome dyads (Figure 8), suggesting a correlation between chromatin accessibility and triplex formation. These results which agree with previous findings by Maldonado et al ^73^, showed that triplexes could form away from the dyad, at the entry-exit site of the nucleosome, helping to fix the nucleosome array and in the case of very long lncRNA helping to approach in the space distant regions as suggested by Marti and co-workers ^46^.

**Figure 8.**
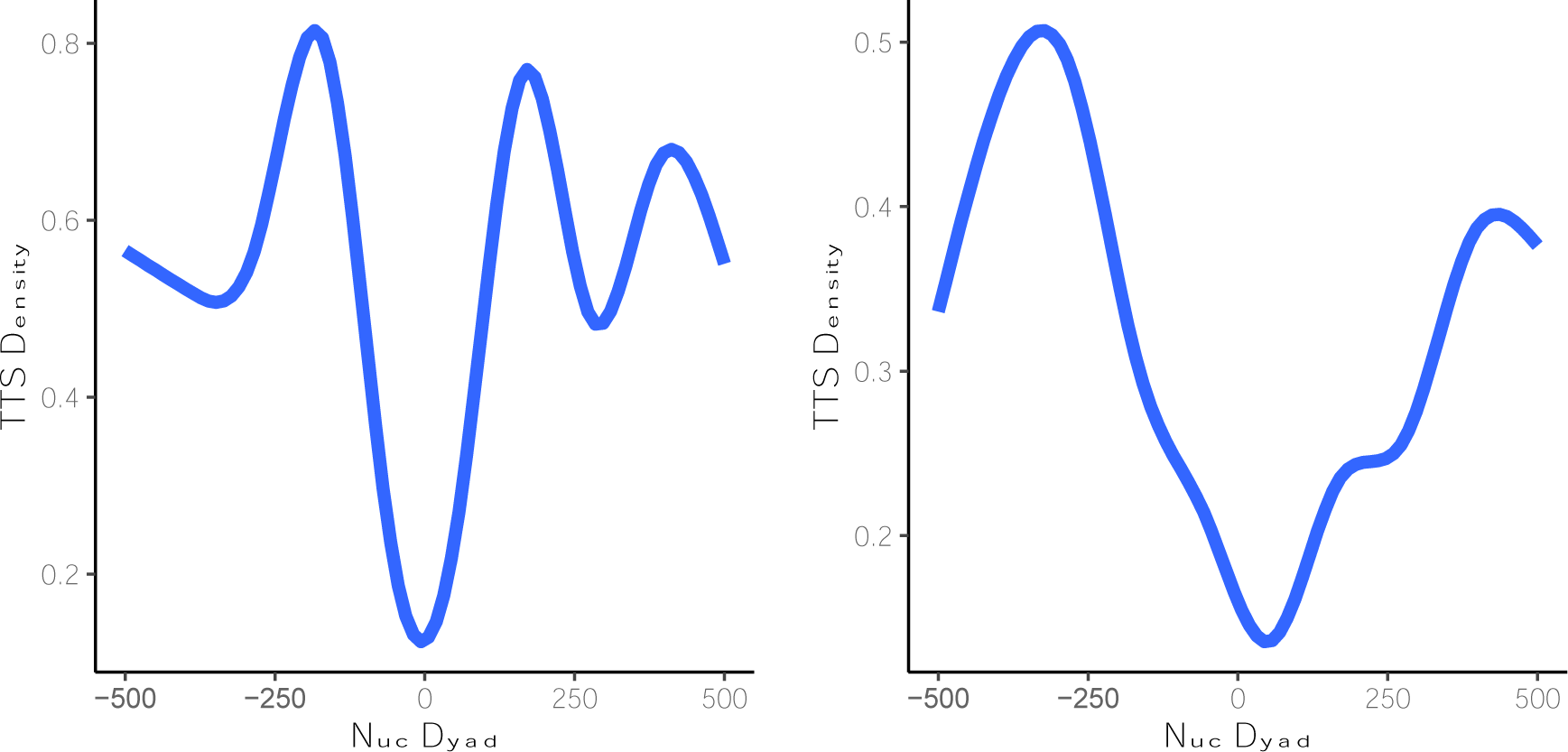
TTS densities from candidate TFOs centered at nucleosome dyads for lncRNAs (left) and miRNAs (right). The nucleosome maps are obtained from lymphoblastoid cells ^72^ and TFOs are originated from lncRNAs and miRNAs expressed in lymphoblastic cells.

## DISCUSSION

A variety of parallel triplexes can be formed mixing complementary DNA and RNA strands, and a significant number of them can be stable under physiological conditions as predicted by state-of-the-art atomistic MD simulations and confirmed by melting and NMR experiments. In general, r(Py)-d(Pu)·r(Py) (*i.e.*, a poly-pyrimidine RNA as TFO and a hybrid DNA(Pur)·RNA(Pyr) as TTS) leads to the most stable structures, followed very closely by the r(Py)-d(Pu)·d(Py) triplex. The formation of the first triplex requires strand invasion of the DNA duplex, and the exposure of an unpaired pyrimidine-rich DNA strand, which would be disfavored in the cell. On the contrary, the r(Py)-d(Pu)·d(Py) triplex can be easily formed without the need for disruption of the DNA duplex, taking as TFO an expressed RNA sequence complementary with the purine strand of the duplex. A massive experimental effort allowed us to develop the first predictor for hybrid triplexes, which despite its simplicity, show quantitative accuracy. This predictor was used to determine all the potential triplexes in human lnc and miRNAs (TFO: expressed RNAs and TTS: genomic DNA). Calculations show a very large number of possible stable triplexes, much more than those predicted by random models. Potential triplexes are concentrated in regulatory regions and UTRs, quite interestingly in genes that are related to the development, morphology and functioning of central nervous system, suggesting a potential role of triplexes in a RNA**←→** DNA mediated regulatory network. This work also suggests that, despite the fact that miRNAs are commonly known as post transcriptional regulators, their nuclear function as transcription regulators via triplex formation is more widespread than at first thought ^74^. Furthermore, mapping potential triplex formation with chromatin structure, we found evidence suggesting a role of triplex formation in fixing nucleosome array probably protecting nucleosome from eviction and in the case of lncRNA helping, as suggested by others, to compact chromatin.

## MATERIALS AND METHODS

### Oligonucleotide Synthesis and Melting Experiments

Hairpins **I-XII** were synthesized as previously described ^63^. Oligonucleotides **XIII** and **7-9** (Figure 7) were synthesized via solid phase synthesis using standard phosphoramidite methods (see Suppl. Methods for details).

Samples containing the required strands were heated to 90 °C and slowly cooled down to allow triplex formation in suitable buffers (see Suppl. Method for details). Melting experiments were performed by heating from low temperature to 100 °C at 0.5 °C/min, monitoring absorbance at 260 nm every 0.5 °C. Experiments were repeated for 5 µM, 8 µM, 12 µM, 18 µM and 22 µM oligonucleotide concentration to derive melting thermodynamic parameters from Van’t Hoff analysis (see Suppl. Methods for details).

### NMR spectroscopy

NMR spectra of hairpin **VII** were first recorded at a range of temperatures (5-45 °C). Later, the TFO was added and the mixture was heated (95 °C) and cooled down slowly, collecting spectra in the same temperature range. Spectra were acquired in a Bruker spectrometer operating at 600 MHz, equipped with cryoprobe and processed with the TOPSPIN software. Water suppression was achieved by including an excitation sculpting module in the pulse sequence ^75^, see Suppl. Methods for additional details.

### Electrophoretic Mobility Shift Assays to Analyze Triplex Formation

TFO **7** was heated at 65 °C for 10 min to prevent self-aggregation and then quickly cooled on ice. Triplex formation was obtained by incubating the TFO with ^32^P-labeled hairpin DNA **XVII** in a suitable buffer for 6 hours at 35 °C (see Suppl. Methods). Electrophoresis was done in a 15% native polyacrylamide gel at 8V/cm for 16 h at pH 5.5 (see Suppl. Methods for details). The gels were analyzed by phosphor-imaging.

### Chromatin Preparation and gDNA purification

HeLa cells were grown to 90% confluency in T75 flask with Dulbecco’s modified Eagle’s medium (DMEM) supplemented with 10% Fetal Bovine Serum (FBS) and 1% penicillin/streptomycin. Cells were trypsinized and nuclei were isolated using standard procedures (see Suppl. Methods) and lysed to obtain chromatin, which was then subjected to treatment with Proteinase K and DNAse I to yield fragments with an average size >10 Kb (Suppl. Figure S3a) followed by phenol/chloroform extraction and ethanol precipitation (see Suppl. Methods for details).

### *In vitro* triplex Pull-down Assay

Purified genomic DNA was sheared into 200-300 bp fragments by sonicating using a Bioruptor Pico (see Suppl. Methods for details). The resulting DNA mixtures were incubated with biotinylated TFO (**8** or **9**; Figure 7) at two different pH values (pH 7.4 and pH 5.5). TFO-associated DNA was captured by incubation with MyOne Streptavidin DynaBeads followed by treatment with RNase H and elution with suitable buffer (see Suppl. Methods for details).

### Parameterization of the Nearest Neighbor Model for RNA-DNA_2_ Triplex Stability

Following Roberts and Crothers ^50^ we determined the enthalpy of triplex formation by (eq. 1):

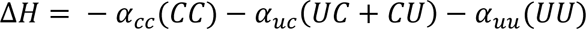

where (XX) refers to the number or dinucleotide steps of the type XpX in the TFO (CC, UC, CU or UU), and αs are fitted parameters.

The ΔG is determined as a function of the nucleotide content and the pH (eq. 2):

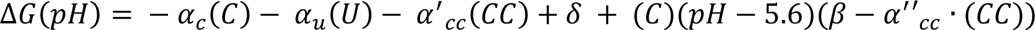

where all symbols in Greek letters are fitted parameters. From ΔH and ΔG, we can extract the T_m_ using (eq. 3) ^76^.

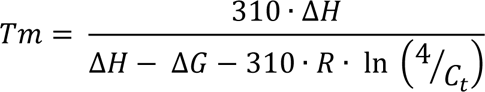

where R is the ideal gas constant and C_t_ is the concentration of the (hairpin) duplex and RNA strands.

The model was parametrized by non-linear fitting using ΔH and ΔG values obtained from our training set (Suppl. Figure S1) at different pH and concentrations (see Suppl. Methods for additional details).

### Bioinformatics Scanning of Potential RNA-DNA·DNA Formation in Humans

We analyzed the triplex potential of annotated lncRNA and miRNA sequences from GENCODE ^68^ and miRbase ^69^ respectively. All annotated sequences were scanned with our stability predictor defining potential TFOs with a minimum length of 10 and a maximum of 30bps. The pH value was set at a default value of 7.0 and the *c_TFO_* was set at a value of 12 μM. A T_m_ of 30°C was set as a threshold to classify stable fragments, while T_m_ of 45°C was considered to detect highly stable triplexes. In order to detect the formation of potential parallel triplex cores (see Suppl. Methods for additional details) we defined our TTSs as polypurine segments with perfect parallel alignment to the previously found TFOs. The extension of the core was evaluated by the melting predictor (see above) with a penalty equal to 10°C decrease in T_m_ per mismatch. The population of potential RNA-DNA·DNA triplexes in the human genome was compared with a random background model. In this model we randomly generated 1 million sequences which followed the base distributions found in the human genome. This allowed us to get a large enough sample for scanning candidate TFOs. We then obtained the target sites from our randomly generated TFOs and used them in the downstream analysis. The potential formation of triplexes in the human genome with our predicted TFOs was validated using the DNA-associated RNA dataset published by Grummt et al. ^45^ and available in GEO repository under the accession number GSE120849.

### Triplex Forming Oligonucleotide Fragment Analysis

Aiming to find the enriched regions where the RNA candidate TFOs would bind, we re-aligned the complementary sequences of our TFOs against the human genome. We used STAR (version 2.5.3a, ^77^) mapping the candidate TTSs to the hg38 assembly of the human genome. The aligned reads were mapped to the corresponding annotation files and classified accordingly. The annotations for genes, exons, transcripts, and UTR regions were obtained from GENCODE (Release 35 (GRCh38.p13)). Promoters were defined as the regions from the transcription start sites up until 1kb upstream. Counts for different features at the gene, promoter regions, exon, transcript and UTR-level resulting from mapping based on the reads were determined separately using the Bioconductor package Rsubread’s function featureCounts (version 2.0.1) ^78^. To further investigate the role of our TTSs, we mapped the obtained promoter sites to their associated genes and performed a Gene Ontology (GO) Analysis using g:Profiler ^79^. A Benjamin-Hochberg FDR index < 0.05 was set to assess significance corrected from multiple test biases. The GO biological processes of the annotated genes were investigated for terms with size > 15 and < 2700 in order to avoid mappings to large pathways that are of limited value and increase statistical value when removing small pathways ^80^.

### Triplex formation in the context of chromatin

The nucleosome map in human lymphoblastoid cell lines was obtained analyzing MNase-seq data (Accession number GSE36979) from Gaffney et al.^81^. The reads were processed with the nucleR package ^82^ as follows: mapped fragments were trimmed to 50 bp maintaining the original center and transformed to reads per million. Noise was filtered through Fast Fourier Transform, keeping 2% of the principal components, and peak calling was performed using the following parameters: peak width 147 bp, peak detection threshold 35%, maximum overlap of 45 bp, dyad length 60 bp. The lncRNA and miRNAs expressed in lymphoblastic cells were obtained from RNA-seq and small RNA-seq data (accession numbers E-MATB-8300 and E-MTAB-8301)^83^.

### Structural Models and Molecular Dynamics (MD) Simulations

Starting conformations for the six triplexes considered here: r(Py)-d(Pu)·r(Py), r(Py)-d(Pu)·d(Py), d(Py)-d(Pu)·d(Py), d(Py)-d(Pu)·r(Py), r(Py)-r(Pu)·r(Py) and r(Py)-r(Pu)·d(Py) were built from DNA triplex structures ^17,18,84,85^. Systems were solvated with waters, neutralized with Na^+^ adding 100 mM additional NaCl. The size of the final triclinic box was approximately 60 Å × 60 Å × 60 Å. Simulation systems were optimized and slowly heated and equilibrated for 50 ns prior to production that extended from 250 ns in the isothermal isobaric ensemble (NPT; T=310 K and P= 1atm). Long-range electrostatic interactions were calculated with the particle mesh Ewald method (PME) with a real space cut-off of 12 Å and periodic boundary conditions in the three directions of Cartesian space were used ^86^. Constant temperature was imposed using Langevin dynamics ^87^ with a damping coefficient of 1 ps, while pressure was maintained with Langevin-Piston dynamics ^88^ with a 200 fs decay period and a 50 fs time constant. LINCS ^89^ was used to maintain covalent bonds at equilibrium distance, allowing the use of 2 fs integration step. Parmbsc1 was used to describe DNA interactions ^64–66^, while RNA was described using recent RNA force-field published by Tan, D. et *al* ^90^. Water molecules were represented by the TIP3P^91^ model, while ions were modeled by Dang’s parameters ^92^.

All MD simulations were performed using *GRO*ningen *MA*chine for *C*hemical *S*imulations (GROMACS) 2016 code ^93^. Analysis of the trajectories were perfomed using GROMACS. Coordinates of the systems were collected every 5 ps of the production trajectory. Analyses were carried out using GROMACS analysis tools, VMD 1.9 Software ^94^, Curves+ ^95^ and NAFlex ^96^ and BIGNAsim analysis tools ^97,98^. Trajectories were stored in our BIGNAsim database ^97,98^ following FAIR data standards as described elsewhere ^98^.

## Acknowledgments

V.G. thanks the European Molecular Biology Organization (EMBO) for financial support (ALTF 103-2018) and “Juan De La Cierva Fellowship” (IJC2019-040468-I / 25A04100). M.O. thanks Spanish Ministry of Science [RTI2018-096704-B-100]; European Research Council (ERC SimDNA), MINECO Severo Ochoa Award of Excellence (Government of Spain) (awarded to IRB Barcelona); the Biomolecular and Bioinformatics Resources Platform (ISCIII PT 13/000/0030 co-funded by the Fondo Europeo de Desarrollo Regional [FEDER]) and the H2020 BioExcel Center of Excellence.

