## Supplementary Information for "Systematic study of hybrid triplex topology and stability suggests a general triplex-mediated regulatory mechanism"

---

[1]Institute for Research in Biomedicine (IRB Barcelona), The Barcelona Institute of Science and Technology, Baldiri Reixac 10-12, E-08028 Barcelona, Spain.
[2]Nostrum Biodiscovery, SL. Barcelona, Spain
[3]Department of Chemistry, University of Cambridge, Cambridge, UK.
[4]Institute for Advanced Chemistry of Catalonia (IQAC), CSIC, Networking Center on Bioengineering, Biomaterials and Nanomedicine (CIBER-BBN), E-08034 Barcelona, Spain
[5] Instituto de Química Física Blas Cabrera. CSIC. E-28006. Madrid
[6] Department of Biochemistry and Biomedicine, University of Barcelona, E-08028 Barcelona, Spain.

### Equally contributing authors

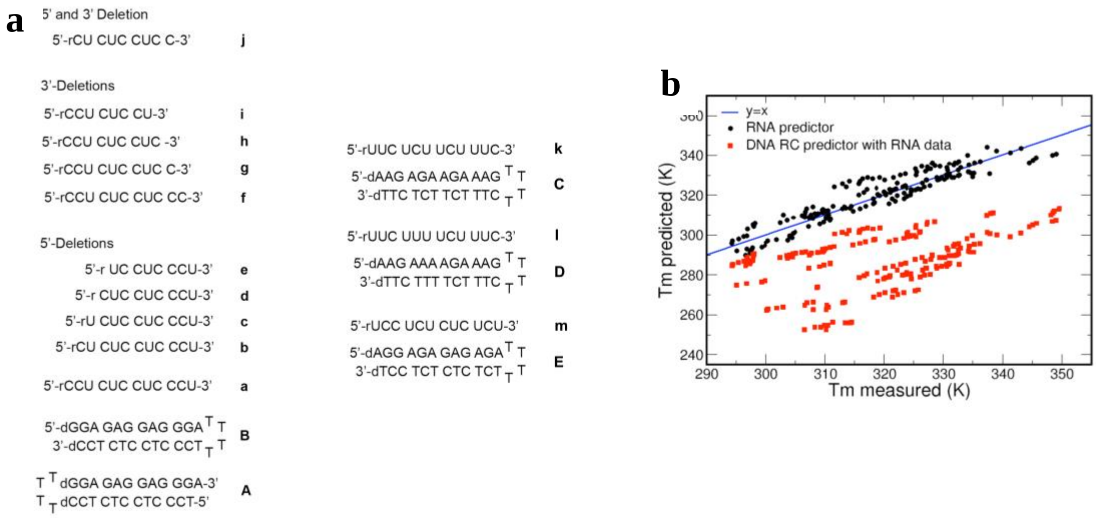


c

| Triplex | $T_m$ (K) Measured | $T_m$ (K) Predicted | ΔH° Measured | ΔH° Predicted | ΔG° Measured | ΔG° Predicted |
|---|---|---|---|---|---|---|
| **A + a** | 335.2 | 336.2 | -91.6 | -83.9 | -17.7 | -16.8 |
| **A + b** | 334.7 | 331.3 | -72.0 | -72.9 | -14.9 | -14.6 |
| **A + c** | 323.3 | 324.2 | -67.1 | -67.2 | -12.6 | -12.7 |
| **A + d** | 321.5 | 322.6 | -61.8 | -61.5 | -12.1 | -12.0 |
| **A + e** | 314.3 | 313.7 | -48.9 | -55.8 | -10.5 | -9.1 |
| **A + f** | 331.0 | 335.7 | -76.2 | -78.2 | -15.2 | -16.1 |
| **A + g** | 330.5 | 330.3 | -73.5 | -67.2 | -14.7 | -13.9 |
| **A + h** | 324.5 | 322.9 | -62.2 | -56.3 | -12.3 | -11.6 |
| **A + i** | 313.0 | 313.3 | -58.6 | -50.5 | -10.4 | -9.7 |
| **A + j** | 330.9 | 330.3 | -69.9 | -67.2 | -14.3 | -13.9 |
| **B + a** | 331.9 | 336.3 | -87.7 | -83.9 | -16.6 | -16.8 |
| **B + b** | 335.1 | 331.3 | -73.7 | -72.9 | -15.6 | -14.6 |
| **B + c** | 326.8 | 324.3 | -65.9 | -67.2 | -13.3 | -12.7 |
| **B + d** | 325.2 | 322.5 | -54.3 | -61.5 | -12.1 | -12.0 |
| **B + e** | 313.5 | 313.7 | -51.8 | -55.8 | -10.2 | -10.1 |
| **B + f** | 328.1 | 335.7 | -74.9 | -78.2 | -14.4 | -16.1 |
| **B + g** | 322.8 | 330.2 | -64.1 | -67.2 | -12.5 | -13.9 |
| **B + h** | 317.1 | 322.9 | -55.3 | -56.3 | -11.0 | -11.6 |
| **B + i** | 307.0 | 313.2 | -44.5 | -50.5 | -9.1 | -9.7 |
| **B + j** | 332.7 | 330.3 | -67.8 | -67.2 | -14.6 | -13.9 |
| **C + k** | 318.2 | 315.8 | -70.8 | -75.9 | -12.1 | -11.0 |
| **D + l** | 306.1 | 309.9 | -64.0 | -67.3 | -9.3 | -9.8 |
| **E + m** | 316.4 | 328.3 | -58.0 | -68.2 | -10.9 | -13.6 |

**Suppl. Figure S1. a.** Sequences used in thermodynamic studies to generate the predictive rPy-dPu·dPy model. **b.** Plot comparing $T_m$ values measured for TFO RNA sequences **a-m** [measured at different pH values (5.0, 5.38, 6.0, 6.5 and 7.0) and oligonucleotide concentration (5 µM, 8 µM, 12 µM, 18 µM and 22 µM)] vs. $T_m$ values predicted for these sequences and conditions using our predictive rPy-dPu·dPy model (black dots) and the dPy-dPu·dPy model developed by Roberts & Crothers (ref 1; red dots). **c.** Comparison of measures and predicted Δ*H*° and Δ*G*° (≈ 37 ºC) for the triplexes shown in panels a-g of this figure, examined at pH 5.0.

| | pH | $T_m$ (K) Measured | Predicted |
|---|---|---|---|
| 5'-rUCU CUC UCU CUC-3' | 5.6 | 335.8 | 329.1 |
| 5'-dAGA GAG AGA GAG $^{T}$ T | 6.5 | 312.7 | 304.3 |
| 3'-dTCT CTC TCT CTC $_{T}$ T | 7.0 | 304.1 | 292.0 |
| 5'-rUCU CCU CUC UCU-3' | 5.6 | 335.2 | 328.3 |
| 5'-dAGA GGA GAG AGA $^{T}$ T | 6.5 | 311.0 | 305.3 |
| 3'-dTCT CCT CTC TCT $_{T}$ T | 7.0 | 298.7 | 293.9 |
| 5'-rCCU CUC UCU CUU-3' | 5.6 | 330.9 | 327.9 |
| 5'-dGGA GAG AGA GAA $^{T}$ T | 6.5 | 310.1 | 305.2 |
| 3'-dCCTCTCTCTCTT $_{T}$ T | 7.0 | 299.8 | 293.9 |
| 5'-rCUU UUC UUU CUC-3' | 5.6 | 311.4 | 315.5 |
| 5'-dGAA AAG AAA GAG $^{T}$ T | 6.5 | 297.3 | 300.8 |
| 3'-dCTT TTC TTT CTC $_{T}$ T | 7.0 | 290.4 | 293.2 |
| 5'-rCUC CUC UCC UCC-3' | 5.6 | 336.6 | 337.6 |
| 5'-dGAG GAG AGG AGG $^{T}$ T | 6.5 | 309.2 | 309.5 |
| 3'-dCTC CTC TCC TCC $_{T}$ T | 7.0 | 295.1 | 295.9 |
| 5'-rUCC CUC CCU CUU-3' | 5.6 | 326.5 | 330.9 |
| 5'-dAGG GAG GGA GAA $^{T}$ T | 6.5 | 302.1 | 308.8 |
| 3'-dTCC CTC CCT CTT $_{T}$ T | 7.0 | 291.7 | 297.7 |

| | pH | $T_m$ (K) Measured | Predicted |
|---|---|---|---|
| 5'-rUUC CUU UCC UU-3' | 5.6 | 313.9 | 314.0 |
| 5'-dAAG GAA AGG AA $^{T}$ T | 6.5 | 298.9 | 300.2 |
| 3'-dTTC CTT TCC TT $_{T}$ T | 7.0 | 291.4 | 293.0 |
| 5'-rUUC CUU UCU UU-3' | 5.6 | 307.9 | 308.0 |
| 5'-dAAG GAA AGA AA $^{T}$ T | 6.5 | 295.4 | 297.3 |
| 3'-dTTC CTT TCT TT $_{T}$ T | 7.0 | 291.9 | 291.6 |
| 5'-rCUC UUC UCC UU-3' | 5.6 | 321.1 | 320.0 |
| 5'-dGAG AAG AGG AA $^{T}$ T | 6.5 | 301.8 | 300.4 |
| 3'-dCTC TTC TCC TT $_{T}$ T | 7.0 | nd | nd |
| 5'-rCUU CUU CCU-3' | 5.6 | 312.1 | 308.3 |
| 5'-dGAA GAA GGA $^{T}$ T | 6.5 | 294.4 | 290.4 |
| 3'-dCTT CTT CCT $_{T}$ T | 7.0 | nd | nd |
| 5'-rCUU CUU CCU UUU C-3' | 5.6 | 319.6 | 322.6 |
| 5'-dGAA GAA GGA AAA G $^{T}$ T | 6.5 | 301.6 | 306.2 |
| 3'-dCTT CTT CCT TTT C $_{T}$ T | 7.0 | 292.4 | 297.8 |
| 5'-rCUC CUC CU-3' | 5.6 | 313.5 | 313.2 |
| 5'-dGAG GAG GA $^{T}$ T | 6.5 | 293.4 | 289.5 |
| 3'-dCTC CTC CT $_{T}$ T | 7.0 | nd | nd |
| 5'-rCUC CUC CUC C-3' | 5.6 | 329.4 | 330.3 |
| 5'-dGAG GAG GAG G $^{T}$ T | 6.5 | 301.9 | 302.7 |
| 3'-dCTC CTC CTC C $_{T}$ T | 7.0 | 292.7 | 289.3 |

**Suppl. Figure S2.** Sequences used in thermodynamic studies for cross-validation of our predictive rPy-dPu·dPy model (validation set), measured at different pH values (5.6, 6.5 and 7.0) and at 18 µM oligonucleotide concentration, and plot comparing measured vs. predicted $T_m$ values. Black dots: comparison of measured $T_m$ values vs. $T_m$ values predicted for TFO RNA sequences shown in Suppl. Figure S1 (**a-m**) using our predictive rPy-dPu·dPy model. Red dots: comparison of measured $T_m$ values vs. $T_m$ values predicted for TFO RNA sequences shown in Figure S1 (**a-m**) using the dPy-dPu·dPy triplex developed by Roberts and Crothers (1).

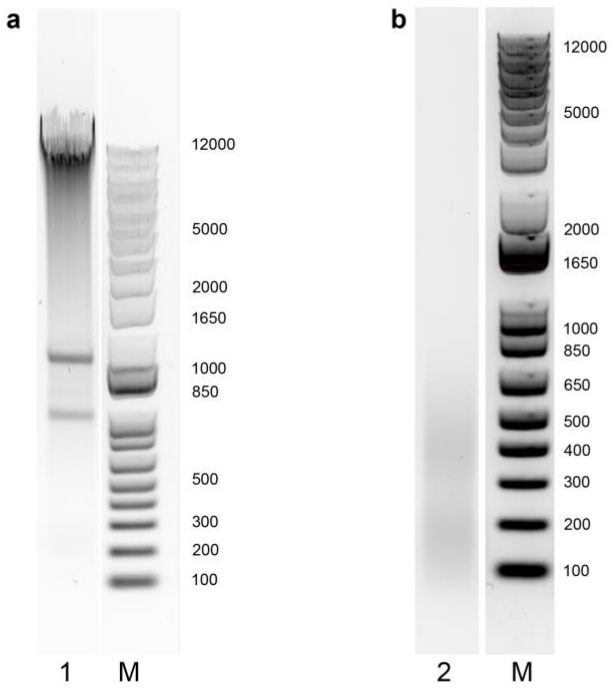


**Suppl. Figure S3. a.** Lane 1: genomic DNA (isolated from HeLa cells) treated (i) with 0.5 unit of DNase I for 5 min at 37 ºC and (ii) with 4 µL of 10% SDS and 4 µL of 20 µg/ µL of Proteinase K for 30 min at 37 ºC. **b.** Lane 2: 10 µg of purified genomic DNA sonicated (6 cycles 30 sec ON/90 sec OFF) in 50 µL of buffer A [10 mM Tris-HCl (pH 7.4), 50 mM KCl, 5 mM $MgCl_2$]. M: DNA ladder.

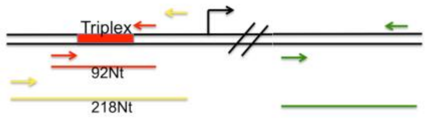


qPCR fragments - triplex capture

For BRD7

Triplex region 92Nt (A) :
GAAAGTGGAAAGAAAGGAAGAGTGGGGGAAGGAGGAAGGGGAAAAAAAGAAAAGAAAAAAAGAGGGAAGGAGACGGGAAGAGGAGAAGGAAG

Triplex region 218Nt (B)
GCTGAGTTTGCTGGGTCTGGCACCCAGTATGGGGTTCAGTCAATATCGTAAAAGGGAAGAAACAGAAGCAAGGGAAAGTGGAAAGAAAGGAAGAGTGGGGGAAGGAGGAAGGGGAA
AAAAAGAAAAGAAAAAAAGAGGGAAGGAGACGGGAAGAGGAGAAGGAAGGGTGATGAGGGAGGAAGAAAAGAGAGAAGGAAGGCTTGAAGGGAGGCAGTAAG

Intron 6 (outside Triplex region) 234 Nt (C)
CTGTTGCTGTGCAGCCCTAAGGAGGTTGCACTCTATTTTACCCAGTGATAACTTTCAGAGTACAGGCAAGGTAGTCAGTAGAAACAAATATGCCTTCTTAAAATTTACGGAGAAACCTCA
AGCCCTTAACTACAATGAAAATTTCACAGCACATCATTGCAGCGAAAAATTAAGCACATGTTTCACAGAATAGAAACAAGACTTCTTTGGTTAGCGAAACCATTCTGCACTCTG

**Suppl. Figure S4.** Schematic representation of the three different regions of BRD7 gene amplified by qPCR.

**a**

| GO:BP | | stats | | | | | |
|---|---|---|---|---|---|---|---|
| Term name | Term ID | $p_{adj}$ | $-\log_{10}(p_{adj})$ | T | Q | T∩Q | U |
| tissue development | GO:0009888 | $1.893\times10^{-6}$ | | 2004 | 526 | 93 | 21010 |
| regulation of developmental process | GO:0050793 | $6.366\times10^{-6}$ | | 2465 | 526 | 106 | 21010 |
| nervous system development | GO:0007399 | $1.499\times10^{-5}$ | | 2513 | 526 | 106 | 21010 |
| neurogenesis | GO:0022008 | $2.741\times10^{-5}$ | | 1708 | 526 | 79 | 21010 |
| skin development | GO:0043588 | $4.691\times10^{-5}$ | | 318 | 526 | 26 | 21010 |
| epidermis development | GO:0008544 | $5.799\times10^{-5}$ | | 388 | 526 | 29 | 21010 |
| epidermal cell differentiation | GO:0009913 | $7.793\times10^{-5}$ | | 245 | 526 | 22 | 21010 |
| keratinocyte differentiation | GO:0030216 | $1.190\times10^{-4}$ | | 175 | 526 | 18 | 21010 |
| regulation of cell differentiation | GO:0045595 | $1.799\times10^{-4}$ | | 1562 | 526 | 71 | 21010 |
| keratinization | GO:0031424 | $2.117\times10^{-4}$ | | 82 | 526 | 12 | 21010 |
| generation of neurons | GO:0048699 | $4.052\times10^{-4}$ | | 1485 | 526 | 67 | 21010 |
| negative regulation of developmental process | GO:0051093 | $6.427\times10^{-4}$ | | 923 | 526 | 47 | 21010 |
| neuron differentiation | GO:0030182 | $1.629\times10^{-3}$ | | 1404 | 526 | 62 | 21010 |
| positive regulation of multicellular organismal proc... | GO:0051240 | $1.911\times10^{-3}$ | | 1629 | 526 | 69 | 21010 |
| regulation of multicellular organismal development | GO:2000026 | $2.508\times10^{-3}$ | | 1396 | 526 | 61 | 21010 |
| potassium ion transmembrane transport | GO:0071805 | $4.807\times10^{-3}$ | | 215 | 526 | 17 | 21010 |
| potassium ion transport | GO:0006813 | $5.277\times10^{-3}$ | | 239 | 526 | 18 | 21010 |
| negative regulation of multicellular organismal pro... | GO:0051241 | $1.062\times10^{-2}$ | | 1102 | 526 | 49 | 21010 |
| adaptive immune response based on somatic reco... | GO:0002460 | $1.158\times10^{-2}$ | | 302 | 526 | 20 | 21010 |
| epithelium development | GO:0060429 | $1.223\times10^{-2}$ | | 1233 | 526 | 53 | 21010 |

**b**

| GO:BP | | stats | | | | | |
|---|---|---|---|---|---|---|---|
| Term name | Term ID | $p_{adj}$ | $-\log_{10}(p_{adj})$ | T | Q | T∩Q | U |
| small molecule metabolic process | GO:0044281 | $1.494\times10^{-23}$ | | 1846 | 2061 | 321 | 21010 |
| cellular response to chemical stimulus | GO:0070887 | $8.235\times10^{-21}$ | | 2691 | 2061 | 416 | 21010 |
| response to external stimulus | GO:0009605 | $1.002\times10^{-20}$ | | 2686 | 2061 | 415 | 21010 |
| regulation of cell population proliferation | GO:0042127 | $1.002\times10^{-20}$ | | 2184 | 2061 | 354 | 21010 |
| positive regulation of multicellular organismal proc... | GO:0051240 | $2.045\times10^{-20}$ | | 1629 | 2061 | 283 | 21010 |
| regulation of developmental process | GO:0050793 | $2.580\times10^{-19}$ | | 2465 | 2061 | 383 | 21010 |
| homeostatic process | GO:0042592 | $2.043\times10^{-18}$ | | 1704 | 2061 | 286 | 21010 |
| tissue development | GO:0009888 | $5.240\times10^{-18}$ | | 2004 | 2061 | 322 | 21010 |
| response to oxygen-containing compound | GO:1901700 | $6.638\times10^{-18}$ | | 1774 | 2061 | 293 | 21010 |
| phosphate-containing compound metabolic process | GO:0006796 | $2.748\times10^{-16}$ | | 2694 | 2061 | 398 | 21010 |
| regulation of immune system process | GO:0002682 | $2.706\times10^{-15}$ | | 1482 | 2061 | 247 | 21010 |
| anatomical structure morphogenesis | GO:0009653 | $5.442\times10^{-15}$ | | 2683 | 2061 | 391 | 21010 |
| regulation of molecular function | GO:0065009 | $1.293\times10^{-14}$ | | 2526 | 2061 | 371 | 21010 |
| regulation of multicellular organismal development | GO:2000026 | $1.886\times10^{-14}$ | | 1396 | 2061 | 233 | 21010 |
| cell adhesion | GO:0007155 | $2.788\times10^{-14}$ | | 1504 | 2061 | 246 | 21010 |
| regulation of localization | GO:0032879 | $4.057\times10^{-14}$ | | 2123 | 2061 | 321 | 21010 |
| response to endogenous stimulus | GO:0009719 | $4.575\times10^{-14}$ | | 1696 | 2061 | 269 | 21010 |
| intracellular signal transduction | GO:0035556 | $2.164\times10^{-13}$ | | 2609 | 2061 | 375 | 21010 |
| negative regulation of multicellular organismal pro... | GO:0051241 | $3.193\times10^{-13}$ | | 1102 | 2061 | 191 | 21010 |
| regulation of transport | GO:0051049 | $5.031\times10^{-13}$ | | 1762 | 2061 | 273 | 21010 |

**c**

| GO:BP | | stats | | | | | |
|---|---|---|---|---|---|---|---|
| Term name | Term ID | $p_{adj}$ | $-\log_{10}(p_{adj})$ | T | Q | T∩Q | U |
| regulation of developmental process | GO:0050793 | $1.456\times10^{-7}$ | | 2465 | 806 | 154 | 21010 |
| nervous system development | GO:0007399 | $4.971\times10^{-6}$ | | 2513 | 806 | 150 | 21010 |
| regulation of cell differentiation | GO:0045595 | $5.865\times10^{-6}$ | | 1562 | 806 | 104 | 21010 |
| multicellular organismal-level homeostasis | GO:0048871 | $5.877\times10^{-6}$ | | 816 | 806 | 65 | 21010 |
| positive regulation of multicellular organismal proc... | GO:0051240 | $1.265\times10^{-5}$ | | 1629 | 806 | 106 | 21010 |
| lipid metabolic process | GO:0006629 | $2.232\times10^{-5}$ | | 1406 | 806 | 94 | 21010 |
| homeostatic process | GO:0042592 | $5.788\times10^{-5}$ | | 1704 | 806 | 107 | 21010 |
| anatomical structure morphogenesis | GO:0009653 | $7.888\times10^{-5}$ | | 2683 | 806 | 152 | 21010 |
| cell migration | GO:0016477 | $8.607\times10^{-5}$ | | 1475 | 806 | 95 | 21010 |
| phosphate-containing compound metabolic process | GO:0006796 | $9.185\times10^{-5}$ | | 2694 | 806 | 152 | 21010 |
| regulation of cell population proliferation | GO:0042127 | $1.189\times10^{-4}$ | | 2184 | 806 | 128 | 21010 |
| cell motility | GO:0048870 | $1.189\times10^{-4}$ | | 1676 | 806 | 104 | 21010 |
| small molecule metabolic process | GO:0044281 | $1.232\times10^{-4}$ | | 1846 | 806 | 112 | 21010 |
| regulation of molecular function | GO:0065009 | $1.576\times10^{-4}$ | | 2526 | 806 | 143 | 21010 |
| tissue development | GO:0009888 | $2.472\times10^{-4}$ | | 2004 | 806 | 118 | 21010 |
| negative regulation of cell differentiation | GO:0045596 | $2.633\times10^{-4}$ | | 676 | 806 | 52 | 21010 |
| fatty acid metabolic process | GO:0006631 | $2.660\times10^{-4}$ | | 396 | 806 | 36 | 21010 |
| negative regulation of developmental process | GO:0051093 | $2.737\times10^{-4}$ | | 923 | 806 | 65 | 21010 |
| neurogenesis | GO:0022008 | $3.875\times10^{-4}$ | | 1708 | 806 | 103 | 21010 |
| cellular lipid metabolic process | GO:0044255 | $5.324\times10^{-4}$ | | 1003 | 806 | 68 | 21010 |

**d**

| GO:BP | | stats | | | | | |
|---|---|---|---|---|---|---|---|
| Term name | Term ID | $p_{adj}$ | $-\log_{10}(p_{adj})$ | T | Q | T∩Q | U |
| small molecule metabolic process | GO:0044281 | $5.583\times10^{-21}$ | | 1846 | 560 | 127 | 21010 |
| response to external stimulus | GO:0009605 | $2.467\times10^{-19}$ | | 2686 | 560 | 156 | 21010 |
| cellular response to chemical stimulus | GO:0070887 | $7.290\times10^{-19}$ | | 2691 | 560 | 155 | 21010 |
| phosphate-containing compound metabolic process | GO:0006796 | $7.290\times10^{-19}$ | | 2694 | 560 | 155 | 21010 |
| response to endogenous stimulus | GO:0009719 | $4.821\times10^{-18}$ | | 1696 | 560 | 114 | 21010 |
| blood circulation | GO:0008015 | $3.645\times10^{-17}$ | | 503 | 560 | 56 | 21010 |
| response to oxygen-containing compound | GO:1901700 | $4.922\times10^{-17}$ | | 1774 | 560 | 115 | 21010 |
| positive regulation of multicellular organismal proc... | GO:0051240 | $4.301\times10^{-16}$ | | 1629 | 560 | 107 | 21010 |
| circulatory system process | GO:0003013 | $4.880\times10^{-16}$ | | 589 | 560 | 59 | 21010 |
| regulation of localization | GO:0032879 | $1.887\times10^{-15}$ | | 2123 | 560 | 125 | 21010 |
| oxoacid metabolic process | GO:0043436 | $5.769\times10^{-15}$ | | 970 | 560 | 76 | 21010 |
| carboxylic acid metabolic process | GO:0019752 | $5.769\times10^{-15}$ | | 948 | 560 | 75 | 21010 |
| response to alcohol | GO:0097305 | $6.575\times10^{-15}$ | | 252 | 560 | 37 | 21010 |
| organic acid metabolic process | GO:0006082 | $7.519\times10^{-15}$ | | 976 | 560 | 76 | 21010 |
| blood vessel diameter maintenance | GO:0097746 | $1.789\times10^{-14}$ | | 142 | 560 | 28 | 21010 |
| regulation of tube diameter | GO:0035296 | $1.789\times10^{-14}$ | | 142 | 560 | 28 | 21010 |
| regulation of tube size | GO:0035150 | $2.109\times10^{-14}$ | | 143 | 560 | 28 | 21010 |
| regulation of system process | GO:0044057 | $2.649\times10^{-14}$ | | 551 | 560 | 54 | 21010 |
| cellular response to organic substance | GO:0071310 | $3.323\times10^{-14}$ | | 2001 | 560 | 117 | 21010 |
| negative regulation of multicellular organismal pro... | GO:0051241 | $4.531\times10^{-14}$ | | 1102 | 560 | 80 | 21010 |

**Suppl. Figure S5.** Top 20 Gene Ontology IDs found with g:Profiler (2) from associated TFOs. Analysis is performed considering promoters from our miRNAs **(a)** and lncRNAs **(b)** candidates; UTRs in miRNAs **(c)** and lncRNAs **(d)** candidates. A Benjamin-Hochberg FDR index < 0.05 was set to assess significance. See Methods for details. Columns indicate Term size (T), Query size (Q), Overlap size (TnQ) and Domain size (U).

#### Supplementary Materials and Methods

##### Oligonucleotide synthesis, deprotection and purification

Oligonucleotides (DNA hairpins and RNA TFOs) used to generate and to validate our R*D·D predictor (sequences shown in Figures S1 and S2) were purchased from Sigma Aldrich. Hairpins **I-XII** (Figure 2, main text) were synthesized as previously described (3). TFOs 1-6 (Figure 2; main text) were purchased from Sigma Aldrich. Oligonucleotides **XIII**, **7-9** (Figure 7; main text) were synthesized via solid phase synthesis using standard phosphoramidite methods (4). Commercially available 5'-*O*-DMT-dC$^{Ac}$-, 5'-*O*-DMT-U-3'-succinyl-LCAA-CPG (Link Technologies) and 3'-protected biotin serinol CPG (Glen Research) were used as the solid supports. Phosphoramidite monomers of dA$^{Bz}$, dC$^{Ac}$, dG$^{iBu}$, T (used in the synthesis of oligonucleotide **XIII**) and 2'-*O*-TBDMS-protected phosphoramidite monomers of A$^{Bz}$, C$^{Ac}$, G$^{dmf}$, U (used in the synthesis of oligonucleotides **7-9**), 3'-protected biotin serinol CPG (used in the synthesis of oligonucleotides **8** and **9**), deblocking solution (3% TCA in $CH_2Cl_2$), activator solution (0.3 M 5-benzylthio-1-H-tetrazole in $CH_3CN$), CAP A solution (acetic anhydride/pyridine/THF), CAP B solution (THF/*N*-methylimidazole 84/16) and oxidizing solution (0.02 M iodine in THF/pyridine/water (7:2:1)) were obtained from commercial sources. All oligonucleotides were synthesized in DMT-ON mode.

Deprotection and purification of oligonucleotides **7-9**: After solid-phase synthesis, the solid support was incubated at 55 ºC for 2 h with 1.5 mL of $NH_3$ solution (33%) and 0.5 mL of ethanol. The supernatant was evaporated to dryness and the residue was treated with triethylamine (75 μL) and triethylamine trihydrofloride (60 μL) in DMSO (115 μL) at 65 ºC for 2.5 h. The oligonucleotides were purified using Glen-Pack Cartridges (Glen Research) following manufacturer's instructions.

Deprotection and purification of oligonucleotide **XIII**: After the solid-phase synthesis, the solid support was incubated at 55 ºC for 17 h with 1 mL of $NH_3$ solution (33%). The oligonucleotide was purified using Glen-Pack Cartridges (Glen Research).

#### Melting experiments

Melting curves were acquired on Varian-Cary-100 spectrophotometer equipped with a thermoprogrammer using the following buffers: 100 mM sodium acetate/acetic acid, 1 mM EDTA (experiments at pH 5.0, 5.38, 5.6) or 100 mM sodium cacodylate/cacodylic acid, 1 mM EDTA (experiments at pH 6.5, 7.0) (see Materials and Methods in the main text for details). Experiments were performed in 1 cm (for 5 µM and 8 µM oligonucleotide concentrations) and 1 mm path length quartz cells (for 12 µM, 18 µM and 22 µM oligonucleotide concentrations).

#### Obtaining thermodynamic parameters from UV melting curves

Following the original work of Robert and Crothers (1), we have modelled the equilibrium of triplex formation as arising from three states, Triplex <-> Hairpin + third strand <-> coil. We have used the $T_m$ dependence on the oligonucleotide concentration to compute thermodynamic parameters for a bimolecular equilibrium between two non-self-complementary (e.g. see Marky and Breslauer (5) and eq. 3 in main text). Accordingly, at each concentration (5, 8, 12, 18 and 22 µM) we computed the $T_m$ and the $\Delta H$ by fitting the derivative of the absorbance with respect to the temperature using van't Hoff equation and the bimolecular equilibria between two non-self-complementary sequences (6). For most cases, the melting curves of the duplex and the triplex are well separated, and therefore we used the equilibrium triplex ↔ duplex + third strand. When the triplex and duplex absorbance transitions overlap, which happens at pH $\leq$ 5.38 for sequence with large cytosine and cytosine dinucleotide contents, we used the equilibrium triplex ↔ duplex+oligo ↔ oligo.

For each triplex, we optimized the fit of all absorbance curves at different concentrations simultaneously, imposing that each share the same $\Delta H$. The linear dependence the inverse of the melting temperature and the natural logarithm of the oligonucleotide concentration (RNA hairpin, in our case) was added as constraint for the overall optimization, which was solved using a non-linear least-square fitting. The selection of the boundaries around the $T_m$ for each triplex-duplex transition can have an impact on the final result. We randomly varied these boundaries and after 2000 fitting iterations we selected the fitting that led to the highest overall coefficient of determination both for each individual melting curve and for the overall linear dependence of the melting temperature and the concentration.

**Parameterization of the pH effect on ΔG**

The ΔG component has the pH dependence included: it's a term that is zero at pH 5.6 and has a dependence on the number of C and CC (i.e. the number of cytosines and cytosine dinucleotides in the sequence). We extracted the ΔG at different pH values (see Figure S1) for the triplexes **A** + **a**, **D** + **l** and **C** + **k** shown in Figure S1. For each sequence composition and pH, we extracted the observed ΔG using the same methodology as described in section XXX. We then calculated the differences between the ΔG at a given pH value and the ΔG predicted by our parameterized model at pH 5.6. We finally fitted a and b in the expression,

$$\Delta G - \Delta G_{pH=5.6} = (C)(pG - 5.6)(a - b(CC))$$

using optimization by minimizing the sum of the squared errors. In this expression, a and b are the unknowns and $\Delta G_{pH=5.6}$ is the ΔG observed at pH 5.6. The RMSE of the fitted correction model was 0.5 kcal/mol.

**NMR spectroscopy**

NMR spectra were acquired in 30 mM phosphate (pH 6.0), 100 mM NaCl, 10 mM $MgCl_2$ buffer (9:1 $H_2O/D_2O$). The equimolar concentration of each oligonucleotide (hairpin **III** and TFO **2**) was 0.5 mM. The hairpin was first dissolved in the buffer and NMR spectra were acquired at 5 ºC, 15 ºC, 25 ºC, 35 ºC and 45 ºC. Then, TFO **2** (dry) was resuspended with the hairpin solution. The resulting solution was heated to 95 ºC, allowed to cool slowly to room temperature and stored at 4 ºC until NMR spectra (at 5 ºC, 15 ºC, 25 ºC, 35 ºC and 45 ºC) were acquired. Spectra were acquired in a Bruker spectrometer operating at 600 MHz, equipped with cryoprobe and processed with the TOPSPIN software. Water suppression was achieved by including an excitation sculpting module in the pulse sequence (7).

**Bioinformatics scanning of potential RNA-DNA·DNA formation in humans.**

The small RNA-seq data of lymphoblastic cells were taken from ArrayExpress under accession number E-MATB-8300 (Damien et al.) (8). Sequence information of the human genome was obtained from the UCSC database (version hg38; December 2013) (9). The reads were reversely mapped against the human genome using STAR (version 2.5.3a, Dobin et al.) (10) to detect the formation of potential parallel triplex cores with the following parameters: --runThreadN 10 --outFilterMultimapNmax 20 --quantMode TranscriptomeSAM GeneCounts --outSAMtype BAM SortedByCoordinate --outSAMattributes All.

**Electrophoretic mobility shift assays to analyze triplex formation**

TFO **21** was heated at 65 ºC for 10 min to prevent self-aggregation and then quickly cooled on ice. Triplex formation was initiated by addition of 3 µL of 3X triplex buffer [135 mM Tris-acetate (pH 5.5), and 30 mM $MgCl_2$], 2 µL $^{32}$P-labeled hairpin DNA **XVII**, 2 µL $H_2O$ containing KCl and 2 µL TFO in a final 9 µL reaction volume. To equilibrate triplex formation, the reaction mixture was incubated at 37 ºC for 6 h. Then, 2 µL 50% glycerol solution containing bromophenol blue was added and the sample was directly loaded onto a 15% native polyacrylamide gel, prepared in 50 mM Tris-acetate (pH 5.5) and 10 mM $MgCl_2$ buffer. Electrophoresis was performed at 8 V/cm for 16 h at 4 ºC in 50 mM Tris-acetate (pH 5.5) and 10 mM $MgCl_2$ buffer and the gel was analyzed by phosphorimaging.

**Cell culture**

HeLa cells were grown in Dulbecco's modified Eagle's medium (DMEM) supplemented with 10% Fetal Bovine Serum (FBS) and 1% penicillin/streptomycin.

**Preparation of chromatin and genomic DNA purification**

HeLa cells were grown in a T75 flask to 90% confluency. Cells were trypsinized by treatment with 1 mL of 0.25% Trypsin at 37 ºC for 5 min. Then, 10 mL of cold DMEM were added. The cell suspension was transferred into a 15 mL falcon tube and centrifuged for 5 min at 200 x g and 4 ºC. The pellet was resuspended in 10 mL PBS and centrifuged for 5 min at 200 x g. Then, the pellet was resuspended in 1 mL PBS, transferred to a 1.5 mL LoBind Eppendorf tube and centrifuged at 200 x g for 2 min. The pellet was treated with 400μL of lysis buffer [10 mM Tris·HCl (pH 7.9), 100 mM KCl, 5 mM $MgCl_2$, 0.5% NP-40, 1 mM DTT] and the resulting suspension was gently pipetted up and down 3-5 times to resuspend the cells. The cell lysate was incubated on ice for 5 min. Meanwhile, a LoBind tube with 1 mL of cold sucrose buffer (10 mM Tris pH 7.4, 150 mM NaCl, 24% sucrose) was prepared. The cell lysate was gently overlaid on top of the sucrose buffer and centrifuged at 3500 x g for 10 min. The pellet was rinsed with 1 mL ice cold PBS-EDTA. Isolated nuclei were resuspended in 500 μL of glycerol/urea buffer [25% glycerol, 20 mM Tris-HCl (pH 7.4), 187.5 mM KCl, 0.5 M urea, 0.5% NP-40, 7.5 mM $MgCl_2$], mixed by vortexing 4 s and incubated on ice for 2 min. The lysate was centrifuged at 13,000 x g for 2 min to precipitate the chromatin-RNA complex. The pellet was briefly rinsed with PBS-EDTA, resuspended with 400 μL of DNAse I buffer [10 mM Tris-HCl (pH 7.5), 2.5 mM $MgCl_2$, 0.1 mM $CaCl_2$] and treated with 0.5 μL of 1 U/ μL DNase I at 37 ºC for 5 min. Then, 4 μL of 10% SDS and 4 μL of 20 μg/ μL of Proteinase K were successively added and the resulting mixture was incubated at 37 ºC for 30 min to yield DNA fragments with an average site of >10 Kb (Figure S3a). gDNA was isolated by phenol/chloroform extraction followed by ethanol precipitation and resuspension in 50 μL RNase free water.

#### *In vitro* triplex pull-down assay

In two different 1.5ml Bioruptor pico microtubes, 10 μg of purified genomic DNA were suspended in 50 μL of buffer A [10 mM Tris-HCl (pH 7.4), 50 mM KCl, 5 mM $MgCl_2$] or buffer B [45 mM Tris-acetate (pH 5.5) 10 mM $MgCl_2$] and sonicated (6 cycles 30 sec ON/90 sec OFF) to yield DNA fragments with an average size of 200-300 bp (Fig. S2b). After sonication, both DNA mixtures were combined in an eppendorf tube and incubated with 20 pmol of biotinylated TFO at 4 ºC for 15 h. After incubation with MyOne Streptavidin C1 Dynabeads for 40 min at room temperature, beads were washed once with 2X B&W buffer (10 mM Tris-HCl (pH 7.5) 1 mM EDTA 2 M NaCl). Putative Rloops were digested by a 30mn incubation at 30ºC with 2.5u RNaseH in 50μl final. Beads were washed 2 times more with 2X B&W buffer. TFO-associated DNA were eluted by incubating the beads 30min at 37ºC with 25 ng/μL RNase A + 2.5u/μl RNase T1 in 50μl final.

Recovered DNA was analyzed by qPCR using 3µl of sample in 10µl reaction using the LightCycler 480 SYBR Green I Master Mix (Roche Diagnostics) and following manufacturer's instructions. Results were normalized to input DNA. Primers listed in Table S3 were used to amplify a 92bp or 218bp fragment containing the triplex region in the promoter of BRD7 or a 234bp fragment in BRD7 intron 6, 34kb downstream of the triplex region.

| | MMG width | mMG width | mG width | H-bonds (WC) | H-bonds (H) | Twist | Roll | Inclination |
|---|---|---|---|---|---|---|---|---|
| **r(Py)-d(Pu)·r(Py)** | 6.3 ± 2.1 | 4.6 ± 0.8 | 9.1 ± 0.9 | 18.3 ± 2.1 | 15.3 ± 1.8 | 29.2 ± 3 | 7.9 ± 5.3 | 14.6 ± 6.4 |
| **r(Py)-d(Pu)·d(Py)** | 9.8 ± 1.8 | 5.1 ± 0.9 | 7.8 ± 1.2 | 18.1 ± 1.7 | 15.1 ± 1.4 | 32.1 ± 2.8 | 3.3 ± 4.9 | 5.6 ± 4.2 |
| **d(Py)-d(Pu)·d(Py)** | 9.7 ± 1.7 | 4.7 ± 1.5 | 6.7 ± 1.4 | 17.6 ± 1.7 | 12 ± 1 | 29.8 ± 2.5 | 3.1 ± 4.5 | 6.6 ± 3.9 |
| **d(Py)-d(Pu)·r(Py)** | 8.6 ± 2 | 3.9 ± 1.2 | 9.3 ± 1.1 | 16.5 ± 1.8 | 11.4 ± 1.7 | 31.2 ± 3.3 | 4.4 ± 4.8 | 8.7 ± 4.3 |
| **r(Py)-r(Pu)·r(Py)** | 7.7 ± 1.3 | 6.6 ± 1.1 | 8.3 ± 0.8 | 16.9 ± 1.8 | 6.7 ± 2.3 | 29.7 ± 3.3 | 4.1 ± 3.7 | 8.4 ± 3.6 |
| **r(Py)-r(Pu)·d(Py)** | 8.4 ± 1.7 | 6.1 ± 0.9 | 8.6 ± 1.3 | 16.6 ± 1.8 | 8 ± 1.7 | 29.4 ± 5.3 | 4.6 ± 3.4 | 7.9 ± 3.2 |

**Suppl. Table S1.** Helical descriptors of the 6 triplexes considered here with the associated standard deviations. Maximum number of Watson Crick and Hoogsteen hydrogen bonds are 19 and 16 respectively Averages were done with the last 100 ns for all triplexes except the unstable r(Py)-r(Pu)·r(Py) and r(Py)-r(Pu)·d(Py), where they were obtained with the first 50 ns of trajectory.

| | %puckering Pu (duplex) | %Puckering Py (duplex) | %Puckering (TFO) |
|---|---|---|---|
| **r(Py)-d(Pu)·r(Py)** | 19% 01'n, 64% C2'n, 15% C1'x | 74% C3'n, 23% C4'x | 65% C3'n, 14% 01'n |
| **r(Py)-d(Pu)·d(Py)** | 32% 01'n, 14% C2'n, 50% C1'x | 42% 01'n, 4% C2'n, 53% C1'x | 96% C3'n |
| **d(Py)-d(Pu)·d(Py)** | 42% 01'n, 4% C2'n, 53% C1'x | 50% 01'n, 4% C2'n, 36% C1'x | 35% 01'n, 6% C2'n, 59% C1'x |
| **d(Py)-d(Pu)·r(Py)** | 22% 01'n, 6% C2'n, 66% C1'x | 61% C3'n, 29% C4'x | 41% 01'n, 2% C2'n, 57% C1'x |
| **r(Py)-r(Pu)·r(Py)** | 81% C3'n, 14% C4'x | 76% C3'n, 19% C4'x | 71% C3'n, 29% 01'n |
| **r(Py)-r(Pu)·d(Py)** | 64% C3'n, 29% C4'x | 73% C3'n, 18% C4'x | 68% C3'n, 33% 01'n |

**Suppl. Table S2.** Average puckering distributions for the 3 strands of the 6 triplexes considered here. The index "n" refers to "endo" and "x" to "exo" conformations. Averages were done with the last 100 ns for all triplexes except the unstable r(Py)-r(Pu)·r(Py) and r(Py)-r(Pu)·d(Py), where they were obtained with the first 50 ns of trajectory.

| | Primer name | Sequence | Amplicon |
|---|---|---|---|
| BRD7_92bp | Primer F | GAAAGTGGAAAGAAAGGAAGAGTGG | 92bp |
| | Primer R | CTTCCTTCTCCTCTTCCCGTCTC | |
| BRD7_218bp | Primer F | GCTGAGTTTGCTGGGTCTGG | 218bp |
| | Primer R | CTTACTGCCTCCCTTCAAGCC | |
| BRD7_Intron6_234bp | Primer F | CTGTTGCTGTGCAGCCCTAAG | 234bp |
| | Primer R | CAGAGTGCAGAATGGTTTCGC | |

**Suppl. Table S3.** qPCR primers
